# A novel skeletal-specific adipogenesis pathway defines key origins and adaptations of bone marrow adipocytes with age and disease

**DOI:** 10.1101/2021.01.05.425449

**Authors:** Hero Robles, Xiao Zhang, Kristann L. Magee, Madelyn R. Lorenz, Zhaohua Wang, Charles A. Harris, Clarissa S. Craft, Erica L. Scheller

## Abstract

Bone marrow adipocytes (BMAs) accumulate with age and in diverse disease states. However, their age- and disease-specific origins and adaptations remain unclear, impairing our understanding of their context-specific endocrine functions and relationship with surrounding tissues. In this study, we identified a novel, bone marrow-specific adipogenesis pathway using the *Adipoq*^Cre+/DTA+^ ‘fat free’ mouse (FF), a model in which *Adipoq*-Cre drives diphtheria toxin-induced cell death in all adiponectin-expressing cells. Adiponectin is highly expressed by BMAs, peripheral adipocytes, and a subset of bone marrow stromal progenitor cells with preadipocyte-like characteristics. Consistent with this, FF mice presented with uniform depletion of peripheral white and brown adipose tissues, in addition to loss of BMAs in canonical locations such as the tail vertebrae. However, unexpectedly, a distinctly localized subset of BMAs accumulated with age in FF mice in regions such as the femoral and tibial diaphysis that are generally devoid of bone marrow adipose tissue (BMAT). Ectopic BMAs in FF mice were defined by increased lipid storage and decreased expression of cytokines including hematopoietic support factor *Cxcl12* and adipokines adiponectin, resistin, and adipsin. FF BMAs also displayed resistance to lipolytic stimuli including cold stress and β3-adrenergic agonist CL316,243. This was associated with reduced expression of adrenergic receptors and monoacylglycerol lipase. Global ablation of adiponectin-expressing cells regulated bone accrual in an age- and sex-dependent manner. High bone mass was present early in life and this was more pronounced in females. However, with age, both male and female FF mice had decreased cortical thickness and mineral content. In addition, unlike BMAs in healthy mice, expansion of ectopic BMAs in FF mice was inversely correlated with cortical bone volume fraction. Subcutaneous fat transplant and normalization of systemic metabolic parameters was sufficient to prevent ectopic BMA expansion in FF mice but did not prevent the initial onset of the high bone mass phenotype. Altogether, this defines a novel, secondary adipogenesis pathway that relies on recruitment of adiponectin-negative stromal progenitors. This pathway is unique to the bone marrow and is activated with age and in states of metabolic stress, resulting in expansion of BMAs specialized for lipid storage with compromised lipid mobilization and endocrine function within regions traditionally devoted to hematopoiesis. Our findings further distinguish BMAT from peripheral adipose tissues and contribute to our understanding of BMA origins and adaptation with age and disease.

## INTRODUCTION

Bone marrow adipose tissue (BMAT) is a unique fat depot located within the skeleton. BMAT acts as an endocrine organ and energy storage depot and has the potential to contribute to the regulation of metabolism, hematopoiesis, and bone homeostasis (reviewed in (1)). The development and subsequent regulation of bone marrow adipocytes (BMAs) varies between skeletal sites (2–5) and current work suggests that BMAs are functionally unique within the context of their niche (6). Specifically, the constitutive BMAT (cBMAT) begins to form in distal regions at or slightly before birth, followed by rapid expansion and maturation early in life (3,5). By contrast, the regulated BMAT (rBMAT) develops later and expands with age, generally in areas of active hematopoiesis (3,5). Recent studies in rodents and humans have also highlighted the heterogeneous metabolic properties of BMAs (2,7), suggesting that their capacity for functional support of surrounding cells may change, particularly with age and in states of systemic disease. BMAT expansion occurs in diverse conditions including anorexia, obesity, aging, osteoporosis, hyperlipidemia, estrogen deficiency, and treatment with pharmacotherapies such as glucocorticoids and thiazolidinediones (1,8). Many of these conditions are associated with increased fracture risk. Thus, understanding the context-specific origins and function of BMAs has important implications for development of clinical and pharmacologic strategies to support skeletal and metabolic health.

Genetic causes of lipodystrophy have provided clues about the molecular differences between BMAT and white adipose tissues (WAT) (reviewed in (5)). Congenital generalized lipodystrophy (CGL) is a disorder characterized by complete loss of peripheral adipose tissues and is associated with secondary complications including hypertriglyceridemia, osteosclerosis, insulin resistance, diabetes, and hepatic steatosis (5,9–11). Patients with CGL uniformly lack WAT, however, BMAT is selectively preserved in those with CGL resulting from mutations in *CAV1* (CGL3) or *PTRF* (CGL4) but not *AGPAT2* (CGL1) or *BSCL2* (CGL2) (5). Similarly, all BMAT is retained in *Cav1* knockout mice and cBMAT is present in *Ptrf* knockouts (3). These results in humans and mice suggest that, unlike WAT, BMAT has unique compensatory mechanisms that promote its preservation. In this study, to define the cellular basis for this observation, we examined the formation and regulation of BMAT in the ‘fat free’ *Adipoq*^Cre+/DTA+^ mouse (FF), a novel genetic model of CGL (10,12).

In the FF mouse, any cell that expresses adiponectin (*Adipoq*-Cre+) will express diphtheria toxin A (DTA), leading to DTA-induced cell death (13,14). Adiponectin is a secreted adipokine that is expressed by all brown, white, and BMAT adipocytes in healthy mice, independent of sex (15). Expression of adiponectin also defines the major BMA progenitor, termed ‘Adipo-CAR’ cells (adipogenic CXCL12-abundant reticular) or ‘MALP’ (marrow adipogenic lineage precursor) (16–18). In the FF mouse, we hypothesized that ablation of adiponectin-expressing cells would promote activation of alternate, adiponectin-negative skeletal progenitors to form adipocytes *in vivo* in times of systemic metabolic demand. To test this hypothesis, we performed adiponectin lineage tracing of bone marrow stromal cells and BMAs. We also analyzed age- and sex-associated changes in bone, BMAT, and peripheral adipose tissues in control and FF mice. In addition, we defined the impact of adrenergic stimulation and peripheral fat transplantation on the formation and regulation of BMAT in the FF model. This work refines our understanding of the origins and adaptation of BMAT with age and disease and defines compensatory pathways of adipocyte formation that are unique to the bone marrow and emerge in states of compromised progenitor function and altered lipid load.

## RESULTS

### Adiponectin is expressed by BMAT adipocytes and a subset of stromal progenitor cells

As described previously (15), *Adipoq*^Cre+/mTmG+^ lineage tracing reporter mice were used to localize adiponectin-expressing cell lineages within the skeletal niche. In this model, any cell having expressed adiponectin (*Adipoq*-Cre+) at any time during its genesis will change plasma membrane color from red to green (mT→mG, (19)). Cross-sections of the proximal tibia and tail vertebrae were imaged at 3- and 16-weeks of age in both males and females after immunostaining for green fluorescent protein (GFP), red fluorescent protein (RFP), and perilipin 1 (PLIN1). *Adipoq*^Cre-/mTmG+^ littermates were used as a negative control. In *Adipoq*^Cre+/mTmG+^ male mice, this work confirmed that membrane-localized GFP expression was present in all PLIN1+, rBMAT adipocytes within the proximal tibia (Fig.1A). Prevalent GFP labeling of reticular stromal cells and bone lining cells was also noted (Fig.1A). GFP expression was absent in hematopoietic cells, chondrocytes, and osteocytes (Fig.1A). Similarly, the cells lining the endosteal bone surface were predominantly *GFP/Adipoq* negative (Fig.1A). In negative controls, all cells within the bone, including adipocytes and bone-lining cells, stained positive for RFP and negative for GFP (Fig.1B). Comparable patterns of GFP expression were observed in the proximal tibia of *Adipoq*^Cre+/mTmG+^ female mice at both 3- and 16-weeks of age (Fig.1C and data not shown). In the tail vertebrae, *Adipoq*-Cre traced all PLIN1+ cBMAT adipocytes independent of sex or age, as indicated by GFP (Fig.1D).

**Figure 1.**
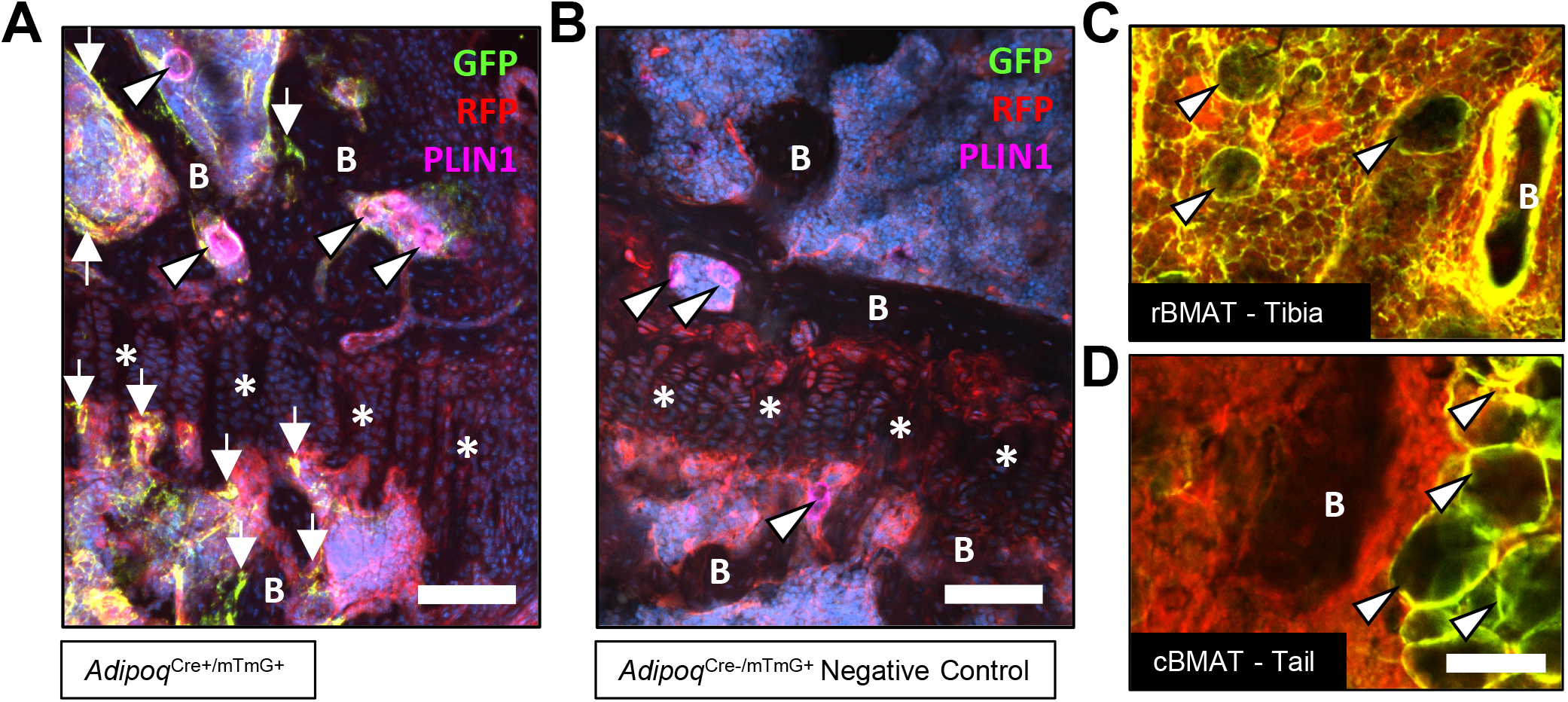
Adiponectin is expressed by BMAT adipocytes in the *Adipoq*^Cre+/mTmG+^ mouse *in vivo*. **(A)** Representative longitudinal cross-section of the proximal tibial metaphysis and epiphysis at the region of the growth plate (*). Bone marrow adipocytes (BMA) were identified by expression of perilipin (PLIN1, pink, arrowheads). BMAs were positive for green fluorescent protein (GFP). GFP expression was also noted in cells lining the bone surface (arrows) and in cells of the stromal reticular network. 4-month old, male *Adipoq*^Cre+/mTmG+^. Scale = 100 μm. **(B)** 4-month old, male *Adipoq*^Cre-/mTmG+^ negative control. All cells, including BMAs, were positive for red fluorescent protein (RFP). Scale = 100 μm. **(C)** Representative proximal tibia, 4-month old, female *Adipoq*^Cre+/mTmG+^. As in males, regulated bone marrow adipose tissue (rBMAT) adipocytes are GFP+. Prominent labeling is also observed in the stromal reticular network and on the bone surface. Scale = 50 μm. **(D)** Representative tail vertebrae, 3-week old male *Adipoq*^Cre+/mTmG+^. Constitutive BMAT adipocytes (cBMAT) are GFP+ (arrowheads). GFP expression is absent in the stroma and on the bone surface. Scale = 50 μm. Images representative of the following animals: 4-month-old *Adipoq*^Cre+/mTmG+^ male (N=5) and female (N=6); 3-week-old *Adipoq*^Cre+/mTmG+^ male (N=5) and female (N=5). B = bone. *Adipoq*^Cre+/mTmG+^ mice were housed at 22°C on a 12h/12h light/dark cycle.

To determine if adiponectin was expressed by the BMAT progenitor cell, we isolated primary bone marrow stromal cells from the femur and tibia of 16-week old male *Adipoq*^Cre+/mTmG+^ mice for colony-forming unit (CFU) assays. After 2-weeks of expansion *ex vivo*, an average of 79.8±9.0% of CFUs were completely positive for adiponectin, as indicated by expression of membrane-bound GFP in 100% of the fibroblast-appearing progenitor cells within the colony, 16.5±9.1% of CFUs were negative (RFP+ only) and 3.7±0.3% were mixed, containing both GFP+ and RFP+ fibroblasts (Fig.2A,B). Within these transitional colonies, cells with RFP+ membranes, indicative of their lack of adiponectin expression, routinely contained GFP+ cytoplasmic granules (Fig.2C). This was often near to cells that had already become fully GFP+ (*Adipoq*-Cre+), suggesting that adiponectin expression is activated at later stages of stromal progenitor maturation. Small, RFP+, myeloid-lineage cells were commonly present, particularly around the edges of the plates (Fig.2A). These cells did not form colonies, did not have a fibroblastic morphology, and thus were not considered in our analyses. Spontaneous adipogenesis, as indicated by the presence of PLIN1+lipid droplets, occurred on average in 18.5+/-1.4% of CFUs (Fig.2D,E). This included 30 of the 166 total colonies examined across three independent mice. Comparable to what was observed in *Adipoq*^Cre+/mTmG+^ mice *in vivo*, PLIN1+ lipid droplets were only present in GFP+ cells *in vitro* (Fig.2D,E). In negative controls, RFP+ stromal cells and PLIN1+ adipocytes were observed without the presence of GFP+ (Fig.2F). Together, these results suggest that all BMAs and their progenitor cells express adiponectin in healthy conditions.

**Figure 2.**
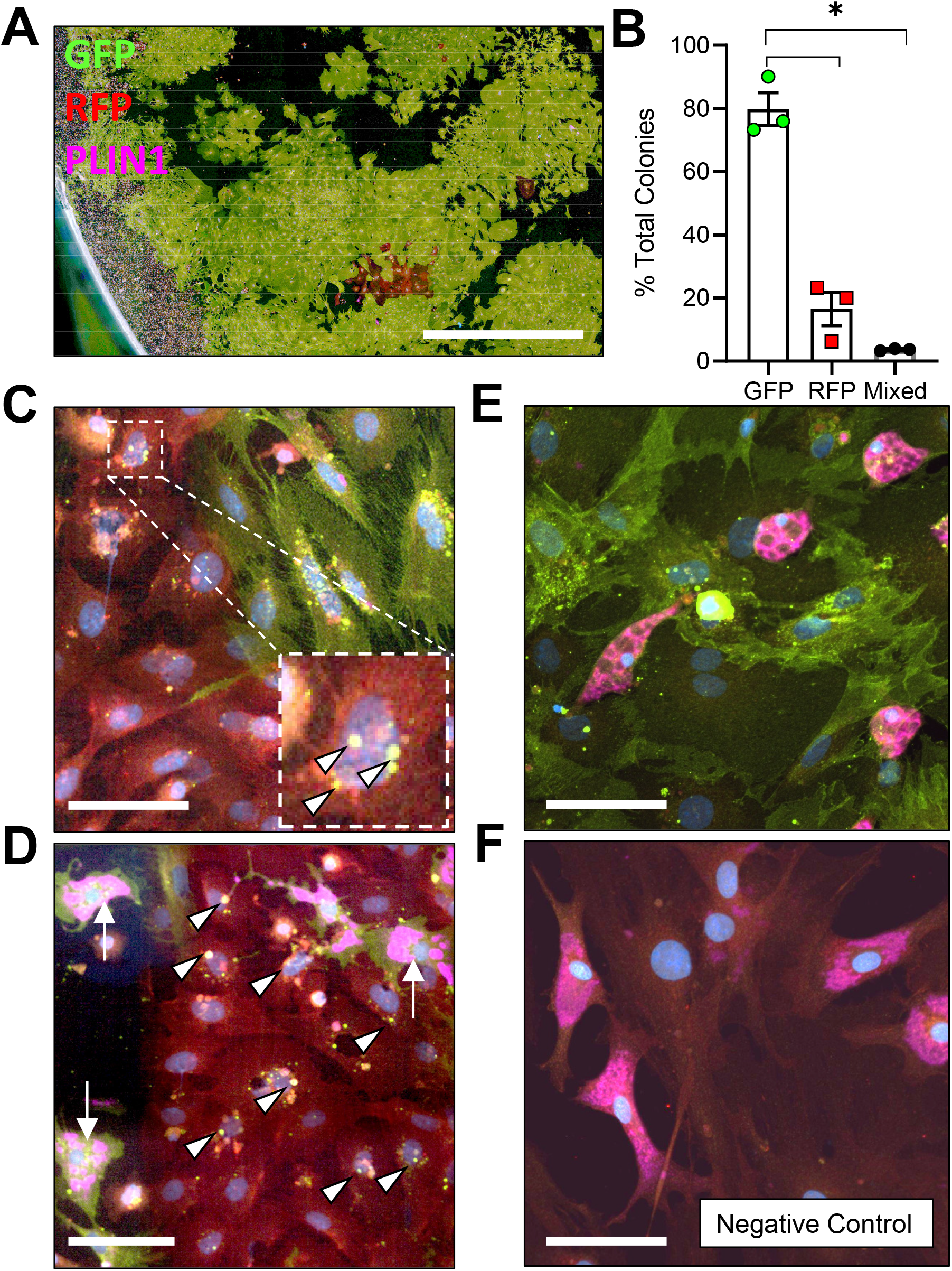
Adiponectin is expressed by *Adipoq*^Cre+/mTmG+^ bone marrow stromal cells *in vitro*. **(A)** Bone marrow stromal cells from 16-week old male *Adipoq*^Cre+/mTmG+^ mice were cultured at low density for 14-days to promote formation of colony-forming units (CFUs). Endogenous fluorescence was then amplified by immunostaining for green fluorescent protein (GFP) and red fluorescent protein (RFP). In addition, spontaneous adipogenesis was assessed based on immunostaining for perilipin 1 (PLIN1, pink). Scale = 1 mm. **(B)** Quantification GFP and RFP expression in fibroblastic stromal cells within each (n = 3 independent mice, 166 total colonies counted). Data presented as mean ± SD. 1-way ANOVA. *p≤0.05. **(C)** Representative mixed colony with both GFP+ and RFP+ fibroblasts demonstrating GFP+ perinuclear granules in red cells (white arrowheads), indicating upregulation of adiponectin expression. **(D)** Day 14 adipogenic colony demonstrating PLIN1+ lipid droplets (pink, white arrows) in GFP+ adipocytes. Nearby RFP+ fibroblasts show early signs of conversion (GFP+ perinuclear granules, white arrowheads). **(E)** Day 14 adipogenic colony with uniformly GFP+ stromal cells and PLIN1+ adipocytes. **(F)** *Adipoq*^Cre-/mTmG+^ negative control has RFP+ bone marrow stromal cells and RFP+, PLIN1+ adipocytes. **(C-F)** Scale = 20 μm.

### Global ablation of adiponectin-expressing cells causes sex- and age-dependent regulation of bone

Male and female FF mice were analyzed at 4-months and 8-months of age relative to *Adipoq*^Cre-/DTA+^ control littermates (Con) to isolate sex and age-related changes in body mass, bone, and bone marrow adiposity after ablation of adiponectin-expressing cells. Male and female FF mice lacked white and brown adipose tissues and circulating adiponectin at both 4- and 8-months of age (Fig.3A-C and data not shown). The absence of fat was accompanied by secondary sequelae including pronounced liver enlargement and steatosis (Fig.3B,D) and elevated blood glucose (Fig.3E). Bone size, as indicated by tibia length, was reduced by 3-7% in FF male and female mice relative to controls (Fig.3F). Body mass was unchanged at 4-months. However, from 4-months to 8-months of age, male FF mice resisted age-associated gains in body mass relative to controls (Fig.3G). By contrast, female FF mice were 9-13% heavier than controls at both ages examined and did not exhibit age-associated restriction (Fig.3G).

**Figure 3.**
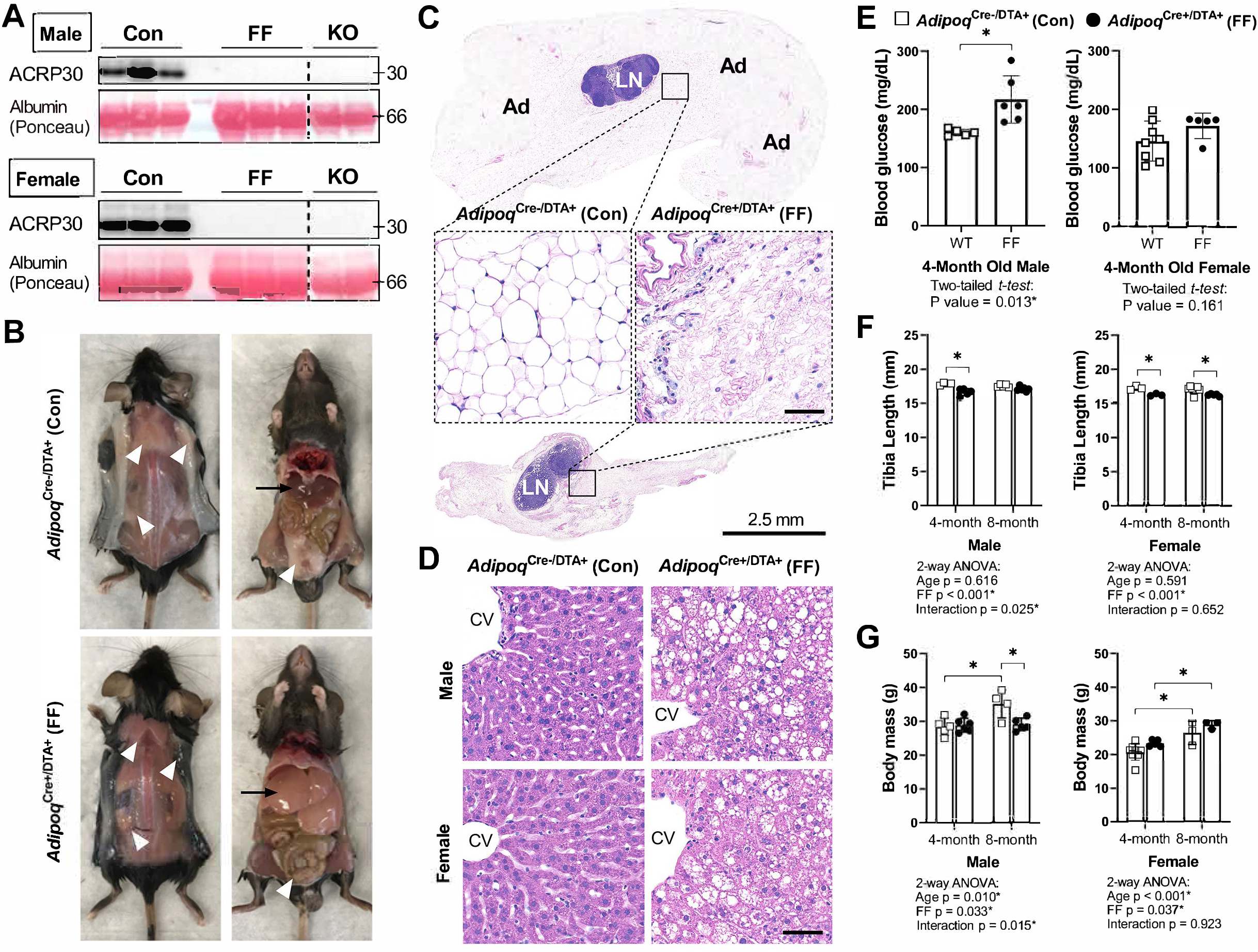
*Adipoq*^Cre+/DTA+^ fat free (FF) mice lack white and brown adipose tissues and circulating adiponectin. **(A)** Serum adiponectin (ACRP30) of *Adipoq*^Cre-/DTA+^ control (Con), *Adipoq*^Cre+/DTA+^ fat free (FF), and *Adipoq* knockout (KO) mice by western blot. Blood albumin levels by Ponceau S staining (loading control). **(B)** Representative pictures showing the absence of white and brown adipose tissues (white arrowheads) and the enlarged liver (black arrows) in 16-week old male FF mice relative to control. Identical gross phenotypes were observed in females (data not shown). **(C)** Representative hematoxylin and eosin (H&E) stained sections of inguinal white adipose tissue. Areas of adipocytes have been replaced by loose fibrous tissue in FF mice. Ad = adipocytes. LN = lymph node. Inset scale = 50 μm. **(D)** Representative (H&E) stained sections of liver. CV = central vein. Scale = 50 μm. **(E)** Random fed blood glucose, measured using a glucometer. **(F)** End point tibia lengths, measured using a caliper. **(G)** Body mass. Sample size for control and FF mice, respectively: 4-months Male n = 5, 6, Female n = 8, 5; and 8-months Male n = 4, 5; Female n = 3, 3. (C) Two-tailed *t-test*, (D,E) 2-way ANOVA with Tukey’s multiple comparisons test. ANOVA results as indicated. *p≤0.05. Data presented as mean ± SD. WT and FF mice were housed at 30°C on a 12h/12h light/dark cycle.

To assess bone morphometry, tibiae were scanned by μCT. Consistent with a previous report in younger males (10), trabecular bone in both male and female FF mice extended deeper into the diaphysis than controls (Fig.4A). In the proximal tibial metaphysis, female FF mice had increased trabecular bone volume fraction (BVF), number, thickness, and bone mineral density (BMD), with decreased trabecular spacing at both 4- and 8-months of age (Fig.4B-F). Increases in metaphyseal trabecular bone were less prominent in the 4-month old male FF mice (Fig.4B). Unlike females, trabecular number was the only factor that was increased significantly in males (Fig.4C) with a comparable decrease in spacing at 4-months of age (Fig.4E). By 8-months of age, metaphyseal trabecular BVF, BMD, number, and spacing in male FF mice were comparable to controls (Fig.4B,C,E,F). In addition, unlike females, male FF mice had decreased trabecular thickness relative to the control group at 8-months of age (Fig.4D). This reveals that ablation of adiponectin-expressing cells is sufficient to promote sustained increases in trabecular bone in females, but not in males.

**Figure 4.**
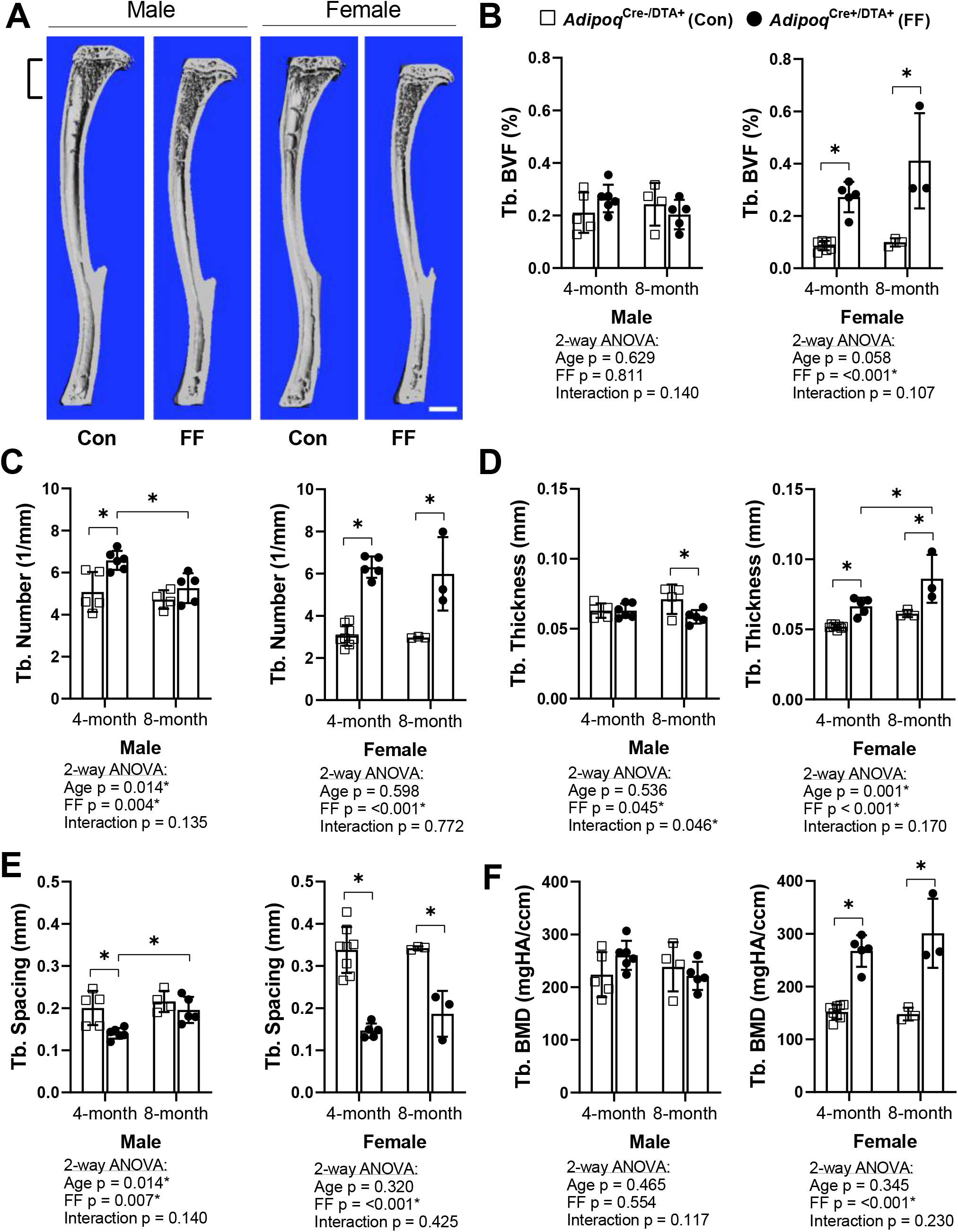
Trabecular bone is increased in *Adipoq*^Cre+/DTA+^ fat free (FF) mice. **(A)** Representative 3D μCT-based reconstructions of tibiae from 4-month old *Adipoq*^Cre+/DTA+^ fat free (FF) and *Adipoq*^Cre-/DTA+^ controls (Con). Scale = 1 mm. **(B-F)** Quantification of trabecular parameters in the proximal tibial metaphysis. Region of interest as indicated in (A). **(B)** Trabecular bone volume fraction (Tb. BVF). **(C)** Trabecular number. **(D)** Trabecular thickness. **(E)** Trabecular spacing. **(F)** Trabecular bone mineral density (Tb. BMD). Sample size for control and FF mice, respectively: 4-months Male n = 5, 6, Female n = 8, 5; and 8-months Male n = 4, 5; Female n = 3, 3. Statistical significance was assessed by two-way ANOVA with Tukey’s multiple comparisons test. ANOVA results as indicated. *p≤0.05. Data presented as mean ± SD. WT and FF mice were housed at 30°C on a 12h/12h light/dark cycle.

Female FF mice at 4-months also had significantly higher cortical BVF and cortical thickness than controls (Fig.S1A-C). Increased cortical thickness was associated with decreased medullary area and no change in total area, indicative of increased endosteal bone (Fig.S1D,E). In male FF mice at 4-months, increased cortical BVF was also associated with decreased medullary area (Fig.S1A-F). However, unlike in females, the total area was also decreased (Fig.S1E), reflecting an overall decrease in bone cross-sectional size. With age, both male and female FF mice exhibited a significant −19.0% and −19.1% decrease in tibial cortical bone thickness, respectively (Fig.S1B). This was in direct contrast to control mice, where tibial cortical thickness remained constant (male) or was increased by +16% (female) with age (Fig.S1B). Changes in the bone mineral content (BMC) mirrored this result, with age-associated increases in controls, but not in FF mice (Fig.S1G). There were no significant differences in predicted torsional bone strength by polar moment of inertia at any of the ages examined (Fig.S1H). Overall, this demonstrates that ablation of adiponectin-expressing cells promotes early gains in the amount and thickness of cortical bone, however, these increases are not sustained and tend to be normalized or decreased relative to controls with age.

### Global ablation of adiponectin-expressing cells drives ectopic expansion of BMAT

Tibiae from the 4- and 8-month old FF and Con mice were decalcified and stained with osmium tetroxide for visualization and quantification of BMAT. Unlike peripheral adipose tissues, the 3D-reconstructed images of the osmium-stained tibiae indicated that BMAT was still present (Fig.5A,B). When quantified and expressed relative to total bone marrow volume, the percentage total tibial BMAT was comparable to controls in 4-month old FF male and female mice and in 8-month old males (Fig.5C). In control mice, BMAT was localized in the well-established pattern of concentration within proximal and distal ends of the tibia (Fig.5A,B) (3). By contrast, the BMAT in the FF mice was found predominantly in the proximal tibia and mid-diaphyseal region with few adipocytes in the distal tibia (Fig.5A,B). Consistent with the 3D reconstructions, regional sub-analyses revealed that retained BMAT adipocytes were primarily localized proximal to the tibia/fibula junction (Fig.5D). Within the proximal tibia, BMAT increased by 2.2-fold in control males and 5.6-fold in control females from 4- to 8-months of age (Fig.5D). In FF mice, though the absolute volume of BMAT was similar or less than controls (Fig.5D), proximal tibial BMAT increased by 6.9-fold and 23.2-fold with age in males and females, respectively (Fig.5D). This included expansion within the mid-diaphysis, a region in mice that is generally relatively devoid of BMAT (Fig.5B). In the distal tibia, control males and females had a large volume of BMAT at 4-months that also increased by 2.7 and 2.3-fold with age (Fig.5E). By contrast, FF mice had very little BMAT in the distal tibia and, though minor increases with age were noted, these changes were not significant (Fig.5E). Distal tibia BMAT often behaves similarly to constitutive BMAT in regions such as the tail vertebrae (2,3). Consistent with this, BMAT adipocytes were generally absent in the 8-month-old FF tail vertebrae, a region of dense cBMAT-like adipocytes in control mice (Fig.5F). These findings demonstrate that BMAT persists in FF mice despite global ablation of adiponectin-expressing cells and, further, that these ectopic BMAT adipocytes expand with age primarily in regions traditionally comprised of red bone marrow.

**Figure 5.**
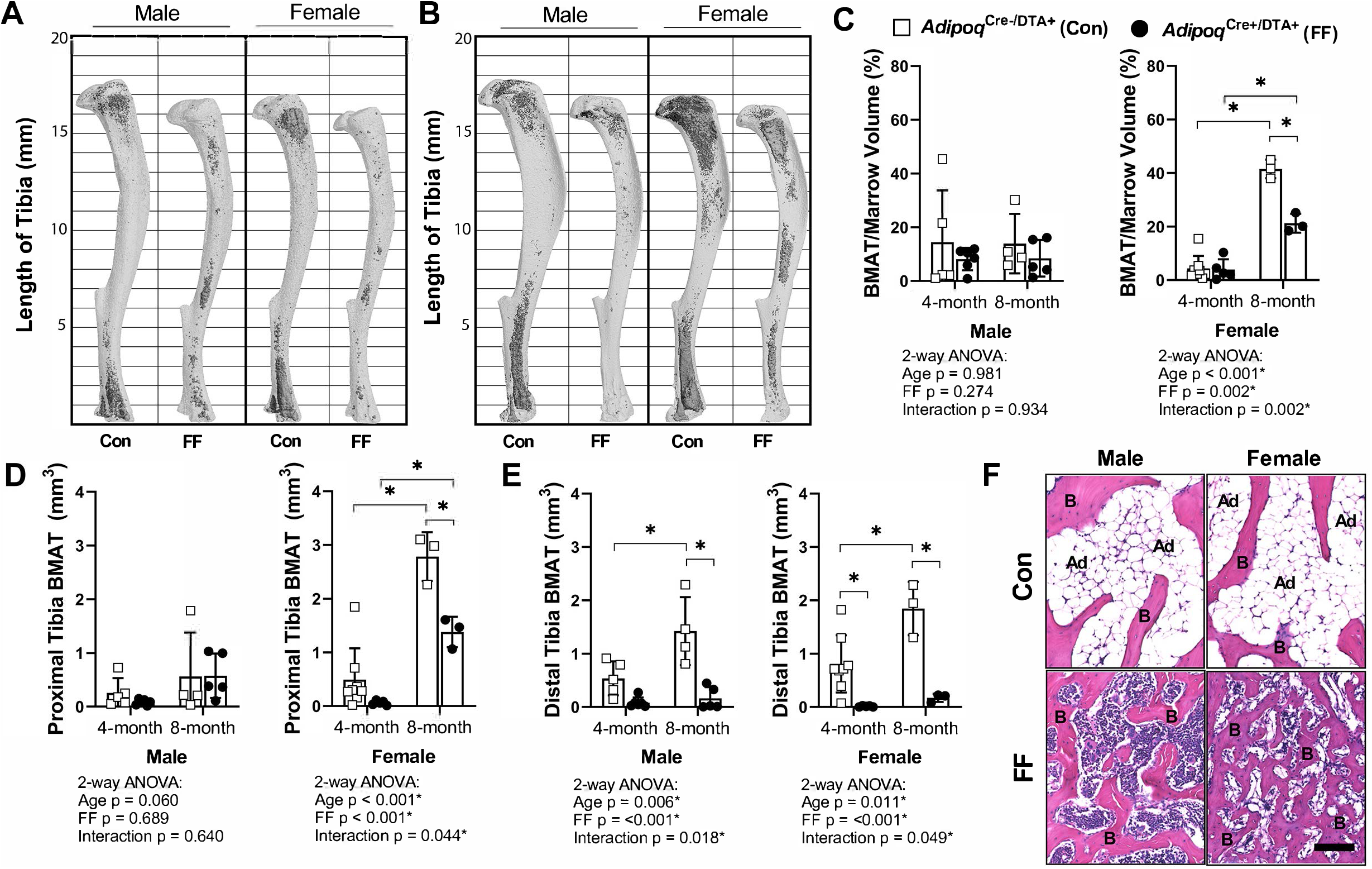
BMAT is present in *Adipoq*^Cre+/DTA+^ fat free (FF) mice and expands with age. **(A,B)** Representative μCT images of osmium-stained tibiae for both male and female *Adipoq*^Cre+/DTA+^ fat free (FF) and *Adipoq*^Cre-/DTA+^ control (Con) mice at **(A)** 4-months and **(B)** 8-months of age. Bone marrow fat is in dark grey and bone is in light grey. **(C)** Quantification of total tibial BMAT volume as a percentage of total bone marrow volume. **(D)** Regional analysis of BMAT within the proximal end of the same tibiae as in (C), expressed as the total volume of osmium-stained lipid from the proximal end of the tibia to the tibia/fibula junction. **(E)** Regional analysis of BMAT within the distal end of the same tibiae as in (C), expressed as the total volume of osmium-stained lipid from tibia/fibula junction to the distal end of the bone. **(F)** Representative hematoxylin and eosin (H&E) stained sections of BMAT within tail vertebrae. Ad = BMAT adipocytes. B = bone. Scale = 50 μm. Sample size for control and FF mice, respectively: 4-months Male n = 5, 6, Female n = 8, 5; and 8-months Male n = 4, 5; Female n = 3, 3. Statistical significance was assessed by two-way ANOVA with Tukey’s multiple comparisons test. ANOVA results as indicated. *p≤0.05. Data presented as mean ± SD. WT and FF mice were housed at 30°C on a 12h/12h light/dark cycle.

By histology, FF BMAT adipocytes in the tibia and femur were morphologically comparable to control BMAT adipocytes (Fig.6A). FF BMAs contained a large, central PLIN1+ lipid droplet (Fig.6B,C) and were negative for macrophage-marker CD68 (Fig.6C). Histologic sections also confirmed the DTA-mediated depletion of the peripheral peri-skeletal adipose tissues in FF mice (Fig.6A). For example, control mice had infrapatellar PLIN1+ adipocytes in the knee joint region (Fig.6A,C). By contrast, in FF mice, the infrapatellar adipocytes were replaced with a population of foam-cell like, auto-fluorescent, PLIN1-, CD68+ macrophages (Fig.6A,C). The same result was observed in the extra-skeletal adipose tissues surrounding the tail vertebrae and the bones in the feet (Fig.S2). This confirms that adipocyte cell death occurs uniformly in the peripheral fat tissues, with selective adipocyte preservation within the bone marrow of FF mice. FF BMAT adipocytes were on average 13.7% and 42.9% larger than controls in male and female mice, respectively, reflecting increases in lipid storage (Fig.6D). Purified FF BMAs also demonstrated a unique gene expression profile. As expected, expression of *Cre* was elevated in FF mice (Fig.6E) with paired decreases in *Adipoq* (Fig.6F). Similarly, expression of cytokines including stromal cell-derived factor 1, also known as C-X-C motif chemokine 12 (*Cxcl12*), adipsin (*Cfd*), and resistin (*Retn*) were significantly decreased (Fig.6F). Expression of adipogenic transcription factor peroxisome proliferator-activated receptor gamma (*Ppary*) was also decreased. By contrast, expression of CCAAT/enhancer-binding protein alpha (*Cebpa*), fatty acid transporter *Cd36*, alkaline phosphatase (*Alpl*), and diphthamide biosynthesis 1 (*Dph1*) were comparable in control and FF BMAs. Overall, this defines the FF BMA as a PLIN+, CD68-adipocyte with increased lipid storage and decreased expression of cytokines including adiponectin, resistin, adipsin, and *Cxcl12*.

**Figure 6.**
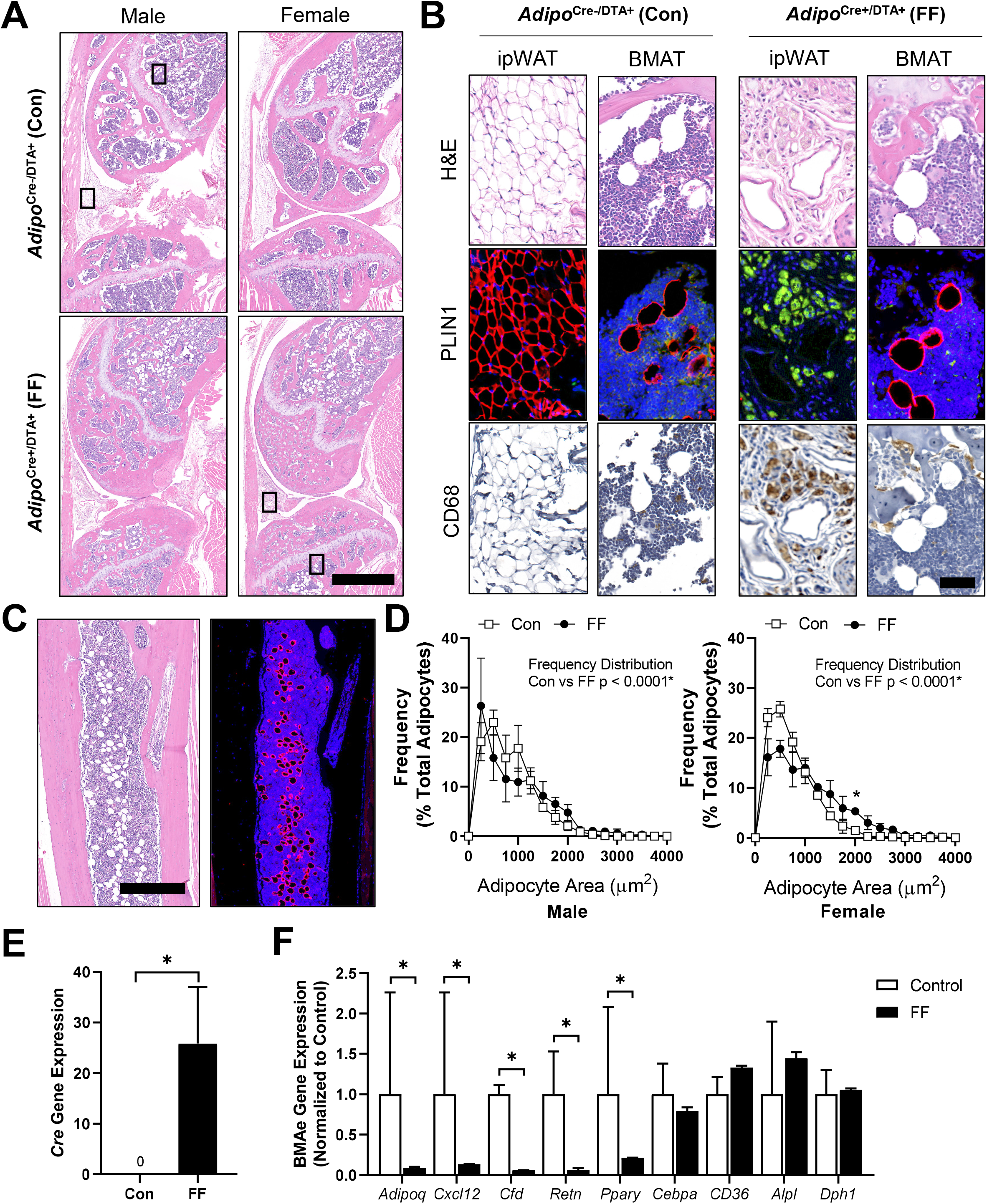
Fat free (FF) bone marrow adipocytes express perilipin, but not CD68, and have increased lipid storage and decreased cytokine expression relative to controls. Representative images from both male and female *Adipoq*^Cre+/DTA+^ fat free (FF) and *Adipoq*^Cre-/DTA+^ control (Con) mice at 8-months of age. **(A)** Representative longitudinal hematoxylin and eosin (H&E) stained sections of the femur and tibia, including the knee and surrounding soft tissue. Scale = 1 mm. **(B)** Representative serial sections stained with H&E, perilipin 1 (PLIN1, red; DAPI, blue), and CD68 (amplified with DAB, CD68+ cells are brown). Sections from the insets depicted in (A). ipWAT = infrapatellar white adipose tissue, located within the knee joint. BMAT = bone marrow adipose tissue. Scale = 50 μm. **(C)** Representative serial sections of the ectopic adipocytes within the tibial diaphysis in the *Adipoq*^Cre+/DTA+^ (DTA) mice. Stained with H&E (left) and perilipin 1 (PLIN1, red; DAPI, blue). **(D)** Bone marrow adipocyte size distribution in the proximal tibia of the control and FF mice at 8-months of age. Scale = 500 μm. Sample size for control and FF mice, respectively: 8-months Male n = 4, 5; Female n = 3, 3. Statistical significance was assessed by two-way ANOVA with Tukey’s multiple comparisons test. ANOVA results as indicated. **(E)** Gene expression of *Cre* normalized to the geometric mean of housekeeping genes *Ppia* and *Tbp* in floated cell preparations enriched for bone marrow adipocytes (BMAe). **(F)** Gene expression in control and FF BMAe preparations, each gene normalized to its respective control. Control n = 2-4, representative of pooled samples from 20-37 mice; FF n = 2, representative of pooled samples from 20 mice. Unpaired t-test with Holm-Sidak correction for multiple comparisons. Data presented as mean ± SD. *p≤0.05. WT and FF mice were housed at 30°C on a 12h/12h light/dark cycle.

### Ectopic BMAT in FF mice is not regulated by cold stress or β3-adrenergic stimulation

Regulation of BMAT adipocytes by adrenergic stimulation has important implications for the functional integration of BMAs with local and peripheral energy stores. FF mice lack WAT and BAT and have impaired thermoregulatory capabilities. Thus, control and FF mice are bred and housed at thermoneutrality (30°C). To assess the response of FF bone and BMAs to thermal stress, male control and FF mice were housed at thermoneutrality (30°C) or room temperature (22°C) for 3-4 months, beginning at 4-weeks of age. Under mild cold stress (22°C), trabecular BVF was decreased by 33-57% relative to housing at thermoneutrality (30°C) in the tibia and femur of control and FF mice (Fig.7A). By comparison, cortical thickness was 9-13% lower in control mice at 22°C but remained unchanged in FF mice, regardless of temperature or analysis site (Fig.7B). By osmium μCT, BMAT in the proximal tibia was 82% lower in control mice housed at 22°C than in mice housed at 30°C (Fig.7C,D). However, similar to cortical bone, proximal tibial BMAT remained unchanged with mild cold stress in FF mice (Fig.7C,D). Retention of BMAT in the FF mice was also prevalent in the femur and presented with the same atypical pattern of accumulation in the mid-diaphysis. Regulation of BMAT within the femur mirrored that observed in the proximal tibia, though it did not reach statistical significance (Fig.7E,F). Unexpectedly, BMAT in both control and FF mice in the distal tibia increased by 1.5- and 5-fold, respectively, in mice housed at 22°C (Fig.7C,D). Overall, this result indicates that in control mice, trabecular bone, cortical bone, and BMAT are decreased in response to mild cold stress. By contrast, only trabecular bone is regulated by thermal stress at 22°C in the FF mice with no observed changes in cortical bone or BMAT.

**Figure 7.**
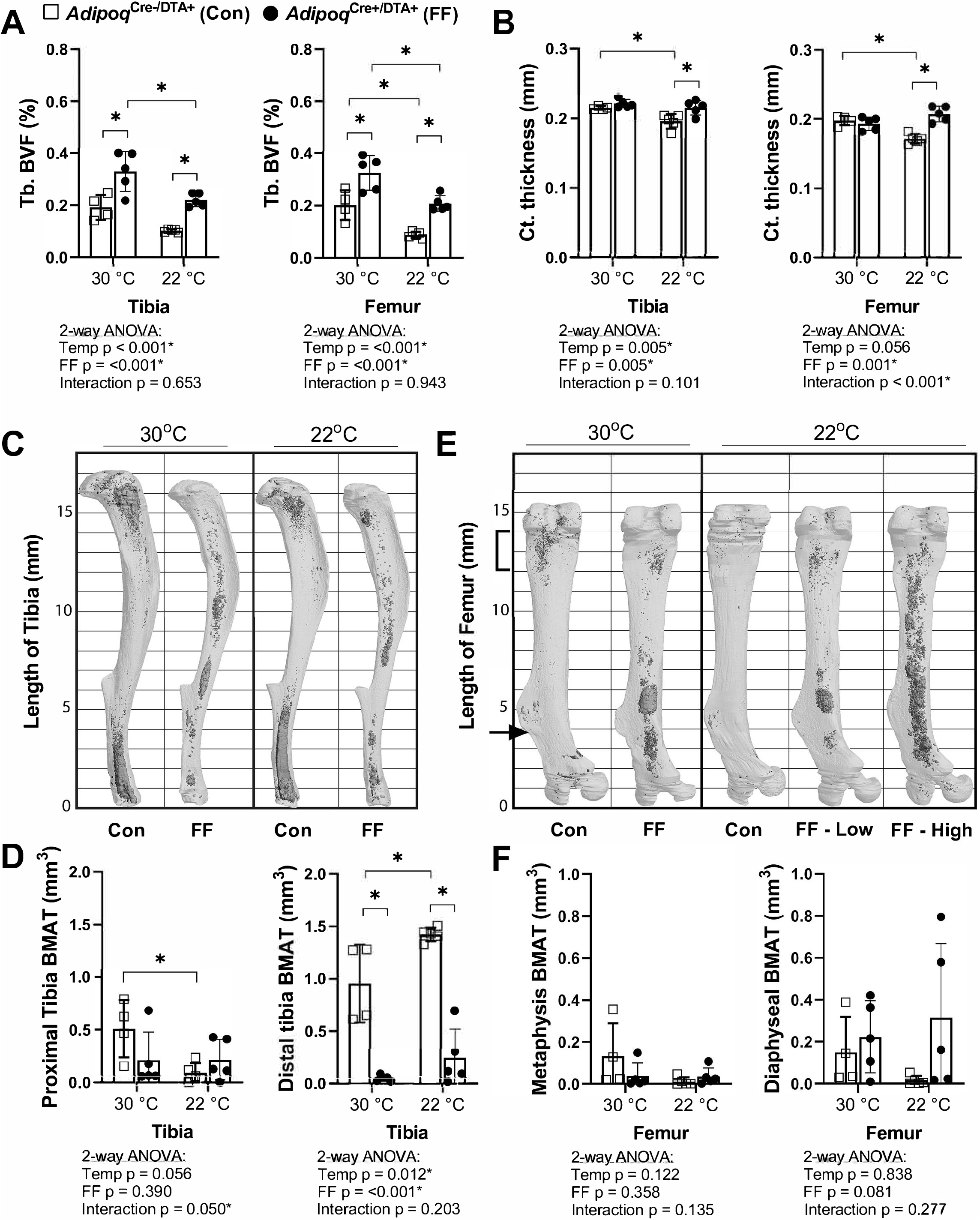
BMAT in *Adipoq*^Cre+/DTA+^ fat free (FF) mice is not responsive to cold temperature challenge (22°C vs 30°C). Male *Adipoq*^Cre+/DTA+^ fat free (FF) mice and controls were maintained in thermoneutral housing (30°C) or moved to room temperature (22°C) at 3- to 5-weeks of age. Bones were analyzed after 3-4 months, at 15- to 17-weeks of age. **(A)** Trabecular bone volume fraction (Tb. BVF) of tibia and femur. **(B)** Cortical thickness of tibia and femur analyzed by μCT. **(C)** Representative μCT images of osmium-stained tibiae at endpoint, respectively. Bone marrow fat is in dark grey and bone is in light grey. **(D)** Quantification of the osmium stained BMAT in the region proximal to the tibia/fibula junction (proximal tibia) or distal to this point (distal tibia). **(E)** Representative μCT images of osmium-stained femur at endpoint, respectively. **(F)** Quantification of the osmium stained BMAT in the 2 mm region below the growth plate (femur metaphysis, bracket) or from this point to the end of the femur flange (indicated by the arrow, diaphyseal BMAT). Sample size of control and FF, respectively: 30°C n = 4, 5; 22°C n = 5, 5. Statistical significance was assessed by two-way ANOVA with Tukey’s multiple comparisons test. ANOVA results as indicated. *p≤0.05. Data presented as mean ± SD.

Next, to isolate responses of control and FF BMAT to direct β-adrenergic stimulation, we treated 7.5-month old male mice with CL316,243, a β3-adrenergic receptor (β3-AR) agonist. Eight daily subcutaneous injections of CL316,243 were administered over the course of 10-days (weekdays only, Monday to Friday in week 1 followed by Monday to Wednesday in week 2) prior to sacrifice on Day 11. To monitor the efficacy of the CL316,243 over time, circulating glycerol concentrations were measured on day 1 and day 7 both immediately prior to and 30-minutes after the CL316,243 injection. Increases in circulating glycerol occur secondary to activation of adipocyte lipolysis and triglyceride hydrolysis by β3-AR (2,20). In control mice, CL316,243 evoked a 2.2-fold and 2.3-fold increase in circulating glycerol on days 1 and 7, respectively (Fig.8A). This response to CL316,243 was absent in FF mice (Fig.8A). After 10-days, β3-AR stimulation decreased BMAT adipocyte cell area in the proximal tibia by 26% in control mice (Fig.9B,C). This reflects an estimated 37% decrease in adipocyte cell volume 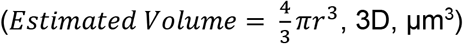. BMAT size was unchanged by β3-AR stimulation in FF mice (Fig.9B,C). Expression of β3 adrenergic receptor (*Adrb3*) was significantly decreased in FF mice (Fig.8D). Gene expression of adipose triglyceride lipase (*Pnpla2*) and hormone-sensitive lipase (*Lipe*) were comparable in FF BMAs (Fig.8D). However, expression of monoglyceride Lipase (*Mgll*) was significantly reduced (Fig.8D). Together, these results suggest that the ectopic BMAT in FF mice is resistant to cold and β3-AR agonist-induced lipolytic stimulation.

**Figure 8.**
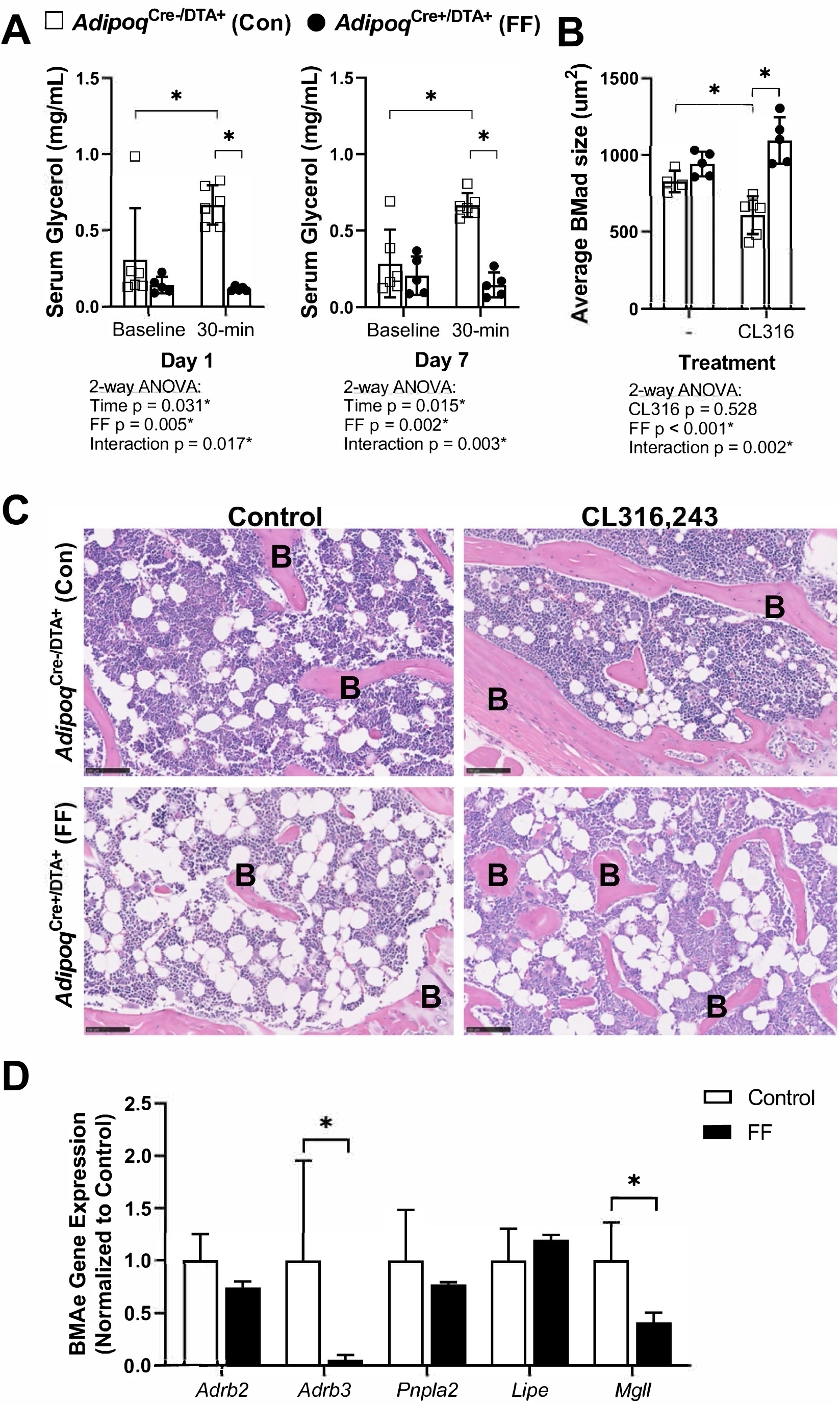
BMAT in *Adipoq*^Cre+/DTA+^ fat free (FF) mice is not regulated by β3-adrenergic stimulation. Male *Adipoq*^Cre+/DTA+^ fat free (FF) and *Adipoq*^Cre-/DTA+^ controls (Con) were treated with CL316,243, a β3-adrenergic receptor (β3-AR) agonist, using a new chronic treatment regimen. Eight daily subcutaneous injections of 0.03 mg/kg CL316,243 were administered to 7.5-month old control and FF mice over the course of 10-days (weekdays only, M→F, M→W) prior to sacrifice on Day 11. **(A)** Serum glycerol at days 1 and 7 of the treatment regimen. **(B)** Average adipocyte cell size in the proximal tibia as assessed in ImageJ using H&E stained slides. **(C)** Representative H&E stained sections. Sample size of control and FF, respectively: 8-month old non-treatment control n = 4, 5 (same mice as in Figs.3–5), 8-month old CL316,243 treated n = 6, 5. Statistical significance was assessed by two-way ANOVA with Tukey’s multiple comparisons test. ANOVA results as indicated. **(D)** Gene expression of the indicated targets normalized to the geometric mean of housekeeping genes *Ppia* and *Tbp* in floated cell preparations enriched for bone marrow adipocytes (BMAe), each gene expressed relative to its respective control. Control n = 2-4, representative of pooled samples from 20-37 mice; FF n = 2, representative of pooled samples from 20 mice. Unpaired t-test with Holm-Sidak correction for multiple comparisons. Data presented as mean ± SD. *p≤0.05. All mice were housed at 30°C on a 12h/12h light/dark cycle.

**Figure 9.**
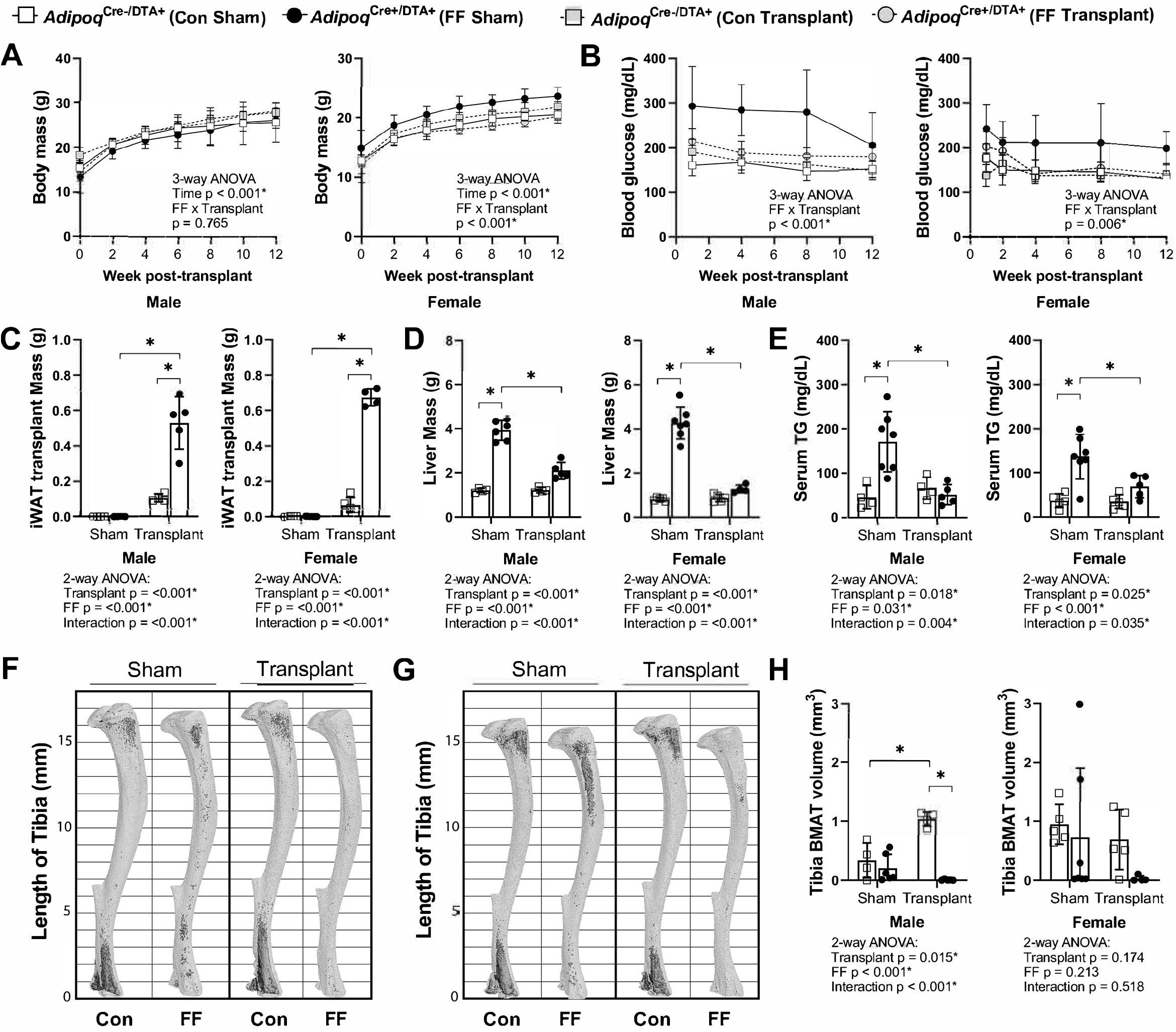
Subcutaneous fat transplant prevents BMAT expansion in *Adipoq*^Cre+/DTA+^ fat free (FF) mice. Male and female *Adipoq*^Cre+/DTA+^ fat free (FF) and *Adipoq*^Cre-/DTA+^ controls (Con) underwent sham surgery or were transplanted subcutaneously with WT inguinal white adipose tissue (iWAT) at 3- to 5-weeks of age. After surgery, mice were monitored for 12-weeks prior to sacrifice. **(A)** Body mass. **(B)** Random fed blood glucose, measured using a glucometer. **(C)** iWAT transplant mass at endpoint (week 12 after transplantation). **(D)** Liver mass at endpoint. **(E)** Serum triglyceride concentration at 4-weeks after the transplant surgery. **(F,G)** Representative μCT images of osmium-stained tibiae of (F) male and (G) female mice at endpoint. Bone marrow fat is in dark grey and bone is in light grey. **(H)** Quantification of total tibial BMAT volume. Sample size for control and FF mice, respectively: sham Male n = 4, 6; transplant Male n = 4, 5; sham Female n = 5, 7; transplant Female n = 5, 4. Statistical significance was assessed by (A,B) three-way ANOVA and (C-H) two-way ANOVA with Tukey’s multiple comparisons test. ANOVA results as indicated. *p≤0.05. Data presented as mean ± SD. All mice were housed at 30°C on a 12h/12h light/dark cycle.

### Subcutaneous fat transplant prevents ectopic BMAT expansion in FF mice

We hypothesized that BMAT expansion in FF mice occurs secondary to peripheral fat depletion and hypertriglyceridemia. To test this hypothesis, male and female FF and control mice underwent sham surgery or were transplanted subcutaneously with wild type adipose tissue at 3- to 5-weeks of age. After surgery, mice were monitored for 12-weeks prior to euthanasia. There were no differences in the body mass of the male mice over time regardless of genotype or transplant (Fig.9A). In females, consistent with previous 4- and 8-month old cohorts (Fig.3E), the body mass of the non-transplanted FF mice was 14-16% higher, on average, than controls (Fig.9A). Subcutaneous fat transplant reduced the body mass of the FF female mice to control levels (Fig.9A). Fat transplant was also sufficient to normalize the hyperglycemia present in both the male and female FF animals (Fig.9B). At the end point, the total mass of the transplanted adipose tissue was significantly higher in the FF mice than in the controls (Fig.9C). As has been reported previously (10), engrafted adipose tissue fragments resembled subcutaneous white adipose tissue at the time of sacrifice (Fig.S3). There was also a significant rescue of liver enlargement and peripheral hypertriglyceridemia by fat transplant in both male and female FF mice (Fig.9D,E). Fat transplant did not substantially modify the cortical and trabecular bone phenotypes in the tibia (Fig.S4). However, the 3D rendering and quantification of tibial BMAT revealed that most of the BMAT present in FF mice was eliminated after subcutaneous fat transplantation (Fig.9F-H). An independent increase in tibial BMAT volume was also observed in male WT fat transplanted mice (Fig.9F,H). The reason for this is unclear as no differences were noted in BAT, iWAT, or gWAT mass after fat transplant in male or female control mice (Fig.S5A-C). Overall, these results reinforce the critical role of the peripheral adipose tissue as a lipid storage depot that reduces the systemic burden of hypertriglyceridemia on peripheral tissues such as liver and bone marrow.

## DISCUSSION

It has previously been assumed that all adipocytes, including bone marrow adipocytes, express the adipokine adiponectin (15,21,22). And, conversely, that adiponectin is not expressed by cells that are not adipocytes. However, recent lineage tracing and single-cell RNAseq studies, including the data presented here, challenge this paradigm and demonstrate that adiponectin is expressed by a subset of bone marrow stromal progenitor cells. These adiponectin-expressing progenitors overlap with CAR cells (17,18,23) and have been more recently termed MALPs (16). They are largely positive for PDGFRβ and V-CAM-1 and have a unique gene expression pattern that mimics known features of pre-adipocytes (17,18,23). Adiponectin-expressing stromal progenitors appear after birth (P1+), matching the known development of BMAT which also occurs primarily postnatally (3,5). Consistent with this and likely due also to the high expression and secretion of adiponectin by healthy BMAs (22), classic depots of rBMAT and cBMAT failed to form in the *Adipoq*^Cre+/DTA+^ FF mouse (3). However, instead, an ectopic population of FF BMAs developed with age in regions of the skeleton such as the diaphysis that are generally devoid of BMAT.

This ectopic BMA population did not rescue circulating adiponectin in the FF mice and had decreased expression of *Adipoq*, reinforcing the efficacy of the DTA. The location of the FF BMAs aligns with known sites of arteriolar entry and distribution within the femur and tibia (24,25). These cells also expressed *Cxcl12*, though this was decreased relative to control BMAs. Arterioles have recently been defined as a site of Osteo-CAR cells, a subpopulation of Cxcl12-expressing cells that are enriched for osteogenic progenitors while Adipo-CAR or MALP cells are primarily localized to the venous sinusoids (16,18). Alternate adiponectin-negative, Cxcl12-negative mesenchymal progenitor populations also exist within bone (26–28). Peri-sinusoidal Adipo-CAR/MALP progenitor cells are generally primed to undergo adipocyte differentiation, however, they are also recruited to undergo differentiation into trabecular bone osteogenic cells with age (~35% at 6-months) and into cortical osteoblasts and osteocytes during injury-induced skeletal repair (17,18) (Fig.10). Our results mirror these findings and support an inverse model whereby the depletion of peri-sinusoidal, adiponectin-expressing MALP/Adipo-CAR progenitor cells drives the preferential differentiation of adiponectin^-/lo^, Cxcl12^-/lo^ progenitors into adipocytes in states of local and systemic metabolic demand, as occurs in CGL (Fig.10). This secondary adipogenesis pathway is unique to the bone marrow and is absent in peripheral adipose tissue depots including WAT and BAT, helping to explain the relative preservation of BMAs relative to WAT in clinical states of CGL and reinforcing their likely importance to maintaining the local homeostasis of the skeletal and hematopoietic microenvironment.

**Figure 10.**
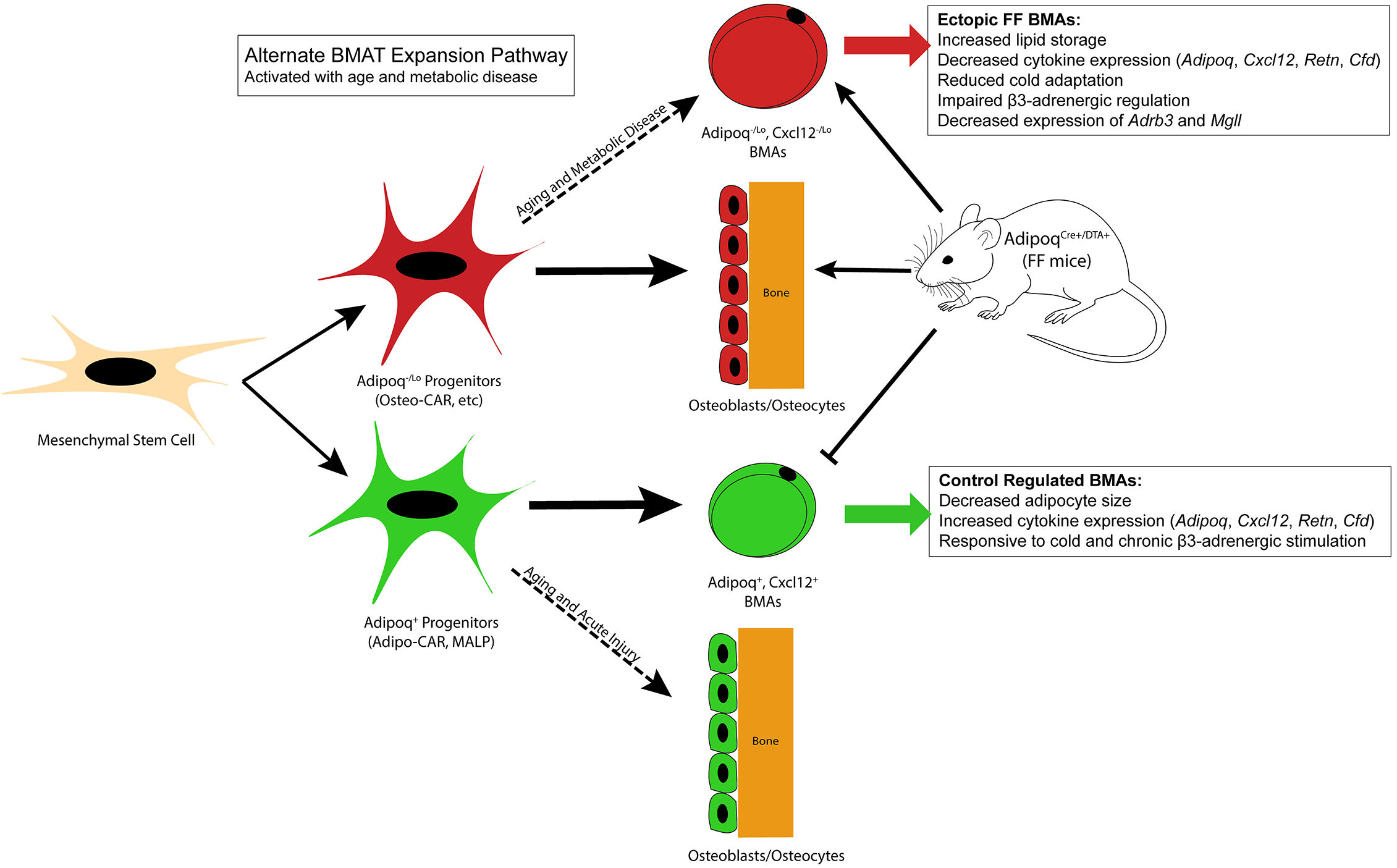
Summary model. Adiponectin is highly expressed by mature bone marrow adipocytes (BMAs) and by a subset of a bone marrow stromal progenitor cells. Adiponectin-expressing progenitors overlap with Cxcl12-abundant reticular (CAR) cells and have been more recently termed Adipo-CAR cells or MALPs (marrow adipogenic lineage precursors). Adipoq+ skeletal progenitors are primed to undergo adipogenesis. Consistent with this and likely due also to the high expression and secretion of adiponectin by healthy BMAs, classic depots of bone marrow adipose tissue (BMAT) failed to form in the *Adipoq*^Cre+/DTA+^ FF mouse. Instead, we observed age-dependent expansion of a BMA population with reduced expression of adiponectin (*Adipoq*^-/lo^) and Cxcl12 (*Cxcl12*^-/lo^) in regions of the skeleton such as the diaphysis that are generally devoid of BMAT. FF BMAs were resistant to cold challenge and β3-adrenergic stimulation and had decreased expression of β3-adrenergic receptors and monoacylglycerol lipase (*Mgll*), suggesting that they have decreased capacity to serve as a local fuel source for surrounding hematopoietic and osteogenic cells. We hypothesize that these cells originate from progenitors including the peri-arteriolar Osteo-CAR population, similar to previous work showing that Adipo-CAR cells are capable of undergoing osteogenic differentiation with age and after acute injury. We propose that expansion of this BMA population is a conserved adaptation with age and in states of metabolic stress and, furthermore, that this is a unique adaptation of the bone marrow that is not present in peripheral adipose depots. Functionally, decreases in stromal and BMA-derived Cxcl12 may contribute to decreased focal support of hematopoiesis, helping to explain the well-defined pattern of bone marrow atrophy and BMA expansion that occurs with age and disease.

The ectopic BMAT in FF mice expands with age and has a larger volume in females than in males, which is a general trend that exists in normal BMAT (29). The 87% decrease in *Cxcl12* expression in FF BMAs also aligns with what has been previously reported in aged BMAs (30). Specifically, a 46% decrease in *Cxcl12* expression was observed in BMAs from 18-month old mice relative to BMAs isolated at 6-months (re-analyzed microarray data from (30)). Decreased BMA-specific expression of *Cxcl12* also occurs in obese mice fed with high-fat diet relative to controls (−24 to −41%, re-analyzed microarray data from (31)). Decreased expression of *Adipoq* has also previously been highlighted as a feature of aged BMAs (30). Thus, we propose that expansion of an adiponectin^-/lo^, Cxcl12^-/lo^ BMA population is a conserved adaptation with age and in states of metabolic stress (Fig.10). Functionally, decreases in stromal and BMA-derived Cxcl12 may contribute to decreased focal support of hematopoiesis (32), helping to explain the well-defined pattern of bone marrow atrophy and BMA expansion that occurs with age and disease (1,5,30). In addition to *Cxcl12* and *Adipoq*, FF BMAs had significant decreases in expression of adipokines including adipsin and resistin, suggesting that these cells may have limited endocrine functions relative to controls.

The deficient response to lipolytic agonists including cold exposure and β3-adrenergic receptor stimulation, larger cell size, and sustained expression of fatty acid transporter *Cd36* also suggests that FF BMAs have decreased capacity to serve as a local fuel source for surrounding hematopoietic and osteogenic cells, preferring instead to take up and to store lipid. This result provides insight into recent conflicting studies on rodent and human BMA lipolysis (2,7). Specifically, purified BMAs from healthy rodents are capable of responding to lipolytic agonists such as forskolin (2). However, purified adipocytes from older humans are not (7). In humans, this was found to be due to a selective decrease in expression of key lipase *Mgll* (7), a serine hydrolase that catalyzes the conversion of monoacylglycerides to free fatty acids and glycerol. Similar to aged human BMAs, we found that FF BMAs have decreased expression of *Mgll* with comparable expression of lipases *Lipe* (Hsl) and *Pnpla2* (Atgl) relative to controls (Fig.8E). This suggests that rodent BMAs can undergo the same adaptations as are present in aged humans, contributing to their resistance to lipolysis. However, unlike humans (7), we also observed decreased expression of *Adrb3* in mice.

Systemic abnormalities including hepatic steatosis, hyperglycemia, and hypertriglyceridemia in FF mice were rescued by subcutaneous fat transplant. Similarly, fat transplant prevented expansion of ectopic BMAs (Fig.9). This supports the existing paradigm whereby excess circulating lipids contribute to the development and expansion of BMAT, and, conversely, that the decrease of these factors in circulation may reduce the development of ectopic BMAs in bone. It is unclear if this could be accomplished in relatively healthy mice or humans to limit age-associated increases in BMAs in regions of hematopoietic bone marrow. And, beyond this, if such a strategy would have benefits to bone or hematopoiesis. The alteration of other circulating factors such as estradiol, leptin, adiponectin, insulin, corticosterone, and catecholamines may also contribute to BMAT expansion in FF mice. Future work is needed to address these questions.

Regarding bone, μCT scans of the FF tibia revealed a general enhancement of bone formation in young FF mice (Fig.4,5), as has been reported previously (10,16). However, this was paired with a mild, but significant decrease in tibial length in FF mice, suggesting that the depletion of adiponectin-expressing cells limits bone elongation, potentially through the suppression of chondrocyte differentiation and/or proliferation within the growth plate. This process may also be mediated by reduced levels of growth hormone or growth factors in circulation or within the bone microenvironment. Consistent with this observation, short stature is consistently reported in patients with CGL3, whose BMAT is also preserved (33). In addition, μCT scans of the FF tibia showed a sex-dependent regulation of bone formation in FF mice. Specifically, trabecular bone formation in FF mice was more pronounced in females with significant differences in age-related trabecular maintenance. It has previously been reported that these *Adipoq*^Cre+/DTA+^ FF mice have low levels of estradiol (34), which may contribute to the sex-dependent changes in bone that were observed in these mice. The onset of this high bone mass phenotype was not altered by peripheral fat transplant or normalization of circulating glucose/lipid levels, supporting previous work by Zhong *et al* and suggesting that changes in bone are driven largely by local depletion of adiponectin-expressing cells (16). Despite early increases in bone, depletion of adiponectin-expressing cells led to suppression of mineral accrual and a decrease in cortical thickness from 4- to 8-months of age in both male and female FF mice, suggesting that short-term gains may give way to longer-term cortical instability. This supports the previous findings of Matsushita *et al* and suggests that adiponectin expressing progenitors, such as MALP/Adipo-CAR cells, may provide necessary contributions to aging or diseased bone through their recruitment to the osteolineage (17).

Historically, BMAT expansion has been linked to bone loss and osteoporosis (8). In recent years, this assumption has come into question as multiple models have shown that both high bone mass and high BMAT can occur simultaneously (3,5,8). In addition, a recent study using *Prx1*-cre mediated knock-out of PPARγ demonstrated that BMAT expansion was not necessary for age-associated bone loss in female mice, though it was sufficient to intensify age-dependent cortical porosity (35). Our study was not designed to isolate the direct relationship between bone and BMAT. However, some observations can be made in the FF model. First, the initial onset of the bone phenotype was independent of the formation of ectopic BMAs, as demonstrated in the fat transplant experiment, suggesting that these early phenotypes are independent. When considered across experiments (n=32 control, 29 FF), trabecular bone volume fraction in the tibia was inversely correlated with metaphyseal BMAT volume in control mice (Slope −2.0, R^2^ 0.137, p=0.037). This is consistent with previous reports in aged mice and humans. However, this association was lost in FF mice (Slope +0.7, R^2^ 0.073, p=0.155). Instead, diaphyseal BMAT was inversely correlated with cortical bone volume fraction in FF mice (Slope −2.1, R^2^ 0.306, p=0.002), but not in controls (Slope +1.8, R^2^ 0.044, p=0.252). In healthy mice, the majority of BMAs are derived from *Adipoq+* marrow adipocyte lineage progenitors (16) (Fig.10). MALP cells form a peri-sinusoidal network throughout the bone marrow and were recently shown to secrete factors such as RANKL, providing active suppression of trabecular, but not cortical, bone mass (36). Ablation of these cells in the FF mice may help to explain the absence of a correlation between FF BMAT and trabecular bone. Instead, this new, ectopic adipocyte population appears to be more closely linked to cortical bone. Though we cannot presume causation, we hypothesize that this relationship may be due to the origination of these BMAs from key adiponectin-negative/low progenitor populations such as Osteo-CAR cells that localize to the peripheral arterioles and endocortical surfaces (18) (Fig.10). Future clarification of this point will undoubtedly help to reveal the niche-specific regulation of and associations between bone, blood, and BMAT.

### Conclusions

With age, BMAs demonstrate gene changes associated with the inflammatory response, mitochondrial dysfunction, and lipid metabolism (30). The unique nature of mature BMA metabolism is not replicated in BMAs differentiated from progenitor cells *in vitro* (7). Consistent with this, our work demonstrates that the spatially defined progenitor cell, the systemic metabolic profile, and the local microenvironment are necessary regulators of BMA expansion and adaptation *in vivo*. In addition, we present evidence for a conserved secondary adipogenesis pathway that is unique to the bone marrow and is activated during times of metabolic stress (Fig.10). The resulting adipocytes express low levels of hematopoietic- and metabolic-supportive cytokines such as *Cxcl12* and *Adipoq*, have increased capacity for lipid storage, and are resistant to lipolytic stimulation, providing new insight into the cellular mechanisms of BMA adaptation. This work refines our understanding of the origins of BMAT and defines compensatory mechanisms of BMAT formation that occur in states of compromised progenitor function and altered lipid load.

## ACKNOWLEDGEMENTS

This work was supported by grants from the National Institutes of Health including R00-DE02417 and startup funds from the Washington University Department of Medicine. We are also grateful for the core services provided by the Musculoskeletal Research Center (NIH P30-AR074992) and the Washington University Center for Cellular Imaging (supported by the Washington University School of Medicine, The Children’s Discovery Institute of Washington University, and St. Louis Children’s Hospital CDI-CORE-2015-505 and CDI-CORE-2019-813 and the Foundation for Barnes-Jewish Hospital 3770 and 4642). Lastly, we would like to extend special thanks to Dr. Jesse Procknow for technical assistance and to Dr. Steven Teitelbaum and Dr. Wei Zou for their helpful discussions during the initial stages of this project.

## AUTHOR CONTRIBUTIONS (based on CRediT taxonomy)

HR – data curation, formal analysis, investigation, methodology, visualization, writing – original draft, writing – review & editing

XZ – conceptualization, data curation, formal analysis, investigation, validation, visualization, writing – original draft, writing – review & editing

KLM – data curation, investigation, writing – review & editing

MRL – data curation, investigation, writing – review & editing

ZW – data curation, investigation, writing – review & editing

CAH – conceptualization, resources, supervision, writing – review & editing

CSC – data curation, formal analysis, funding acquisition, investigation, methodology, visualization, project administration, writing – review & editing

ELS – conceptualization, data curation, formal analysis, funding acquisition, investigation, project administration, resources, supervision, visualization, writing – original draft, writing – review & editing

## METHODS

### Table of key resources

2 Table of key resources
| REAGENT or RESOURCE | SOURCE | IDENTIFIER |
| --- | --- | --- |
| MOUSE MODELS:<br>Background/Strains |  |  |
| C57BL/6J: *Adipoq*-Cre | Jackson Laboratories | 028020 |
| C57BL/6J: ROSA26<sup>mTmG/+</sup> | Jackson Laboratories | 007676 |
| C57BL/6J: ROSA26<sup>DTA/DTA</sup> | Jackson Laboratories | 009669 |
| C57BL/6J: *Adipoq*<sup>-/-</sup> | Jackson Laboratories | 008195 |
| CHEMICALS and KITS |  |  |
| 10% neutral buffered formalin | Fisher Scientific | 23-245684 |
| EDTA | Sigma-Aldrich | E5134 |
| Hydrogen Peroxide | Sigma-Aldrich | 216763 |
| IMMPRESS HRP Anti-Rabbit IgG kit | Vector Laboratories | MP-7401 |
| DAPI | Sigma-Aldrich | D9542 |
| Hematoxylin | Ricca Chemical | 3536-16 |
| OCT mounting media | Fisher HealthCare | 23-730-571 |
| MesenCult™ Expansion Kit (Mouse) | STEMCELL Technologies | 05513 |
| Triton X-100 | Sigma-Aldrich | 9002-93-1 |
| Fluoromount-G | Thermo Fisher Scientific | 00-4958-02 |
| Osmium tetroxide | Electron Microscopy Sciences | 19170 |
| Potassium dichromate | Sigma-Aldrich | 24-4520 |
| Serum Triglyceride Determination Kit | Sigma-Aldrich | TR0100 |
| CL316,243 | Sigma-Aldrich | C5976 |
| Donkey serum | Sigma-Aldrich | D9663 |
| 4x NuPage LDS Buffer | Thermo Fisher Scientific | NP0007 |
| Ponceau S | Fisher Scientific | BP103-10 |
| DNase and protease free bovine serum albumin | Fisher Scientific | BP9706 |
| HBSS Buffer | Gibco | 10425-076 |
| NucleoSpin RNA XS Kit | Takara Biosciences | 740902 |
| SuperScript IV VILO Master Mix with ezDNase™ Enzyme | Thermo Fisher Scientific | 11766050 |
| qPCRBIO SyGreen Mix Lo-ROX | PCR Biosystems | PB20.11-51 |
| Antibodies for immunostaining and western blot | Detailed in Table S1 | N/A |
| SOFTWARE |  |  |
| GraphPad Prism | GraphPad | v8.4.3 |
| SCANCO Medical microCT systems | Scanco Medical AG | N/A |
| NDP.view2 | Hamamatsu Photonics | U12388-01 |
| Fiji | ImageJ | N/A |
| Microsoft Excel | Microsoft | N/A |
| IMAGING SYSTEMS |  |  |
| Scanco µCT 40 | Scanco Medical AG | N/A |
| 2.0-HT NanoZoomer System | Hamamatsu Photonics | N/A |
| Spinning Disk Confocal Microscope | Nikon | N/A |
| LI-COR Odyssey Imager | LI-COR | N/A |
| OTHER |  |  |
| Glucometer | Contour Next | N/A |
| Digital Caliper | iKKEGOL | 5486 |
| PicoLab Rodent Diet 20 | LabDiet | 5053 |
| Microplate Spectrophotometer | BioTek | Epoch |
| QuantStudio™ 3 Real-Time PCR System | Thermo Fisher Scientific A28136 | A28136 |

### Mice

Institutional guidelines for the handling and experimentation with animals were followed and all work was approved by the animal use and care committee at Washington University (Saint Louis, MO, USA). All animals were housed on a 12-hour light/dark cycle and fed *ad libitum* (PicoLab 5053, LabDiet). Mice were obtained from Jackson Laboratories and bred at Washington University including *Adipoq*-Cre (Strain #028020), ROSA26^mTmG/+^ (Strain #007676), ROSA26^DTA/DTA^ (Strain #009669), and Adipoq^-/-^ (Strain #008195). For breeding, heterozygous *Adipoq*-Cre+ males were bred to homozygous ROSA26^mTmG^ or ROSA26^DTA^ females to generate lineage reporter (*Adipoq*^Cre+/mTmG+^) or FF (*Adipoq*^Cre+/DTA+^) mice and associated control littermates (*Adipoq*^Cre-/mTmG+^ or *Adipoq*^Cre-/DTA+^). Mice expressing DTA under the control of the *Adipoq*-Cre promoter lack both white and brown adipose tissues and were bred and housed at thermoneutral temperature (30°C). All transgenic mice were maintained on a C57BL/6J background (Strain #000664). Body mass was recorded with an electronic scale and blood glucose was monitored by tail prick with a glucometer (Contour Next). For end points requiring tissue mass measurements, mice were euthanized with carbon dioxide followed by cervical dislocation. Tissues were collected and weighed using an electronic scale. For end points requiring histology and immunostaining, mice were anesthetized with ketamine/xylazine cocktail (100mg/kg ketamine; 10 mg/kg xyalzine) and perfused through the left ventricle of the heart with 10 mL phosphate-buffered saline followed by 10 mL 10% neutral buffered formalin (NBF, Fisher Scientific 23-245684). When indicated, tibia and femur lengths were determined using a digital caliper (iKKEGOL). For all experiments, collected tissues were post-fixed in 10% NBF for 24-hours. For western blot and serum assays, as detailed below, blood was collected through capillary action from the lateral tail vein and serum was isolated by centrifugation at 1500 x g for 15 minutes after clotting on ice.

### Western blot

Immunoblotting for serum adiponectin (constant volume) was performed as described previously (22). Specifically, serum samples were reduced and denatured in 4X NuPage LDS sample buffer (ThermoFisher, NP0007) containing 1:8 parts β-mercaptoethanol (2 μL serum + 10 μL LDS buffer + 28 μL water). Preparations were incubated at 95°C for 5-minutes and cooled on ice for 1-minute before separating by SDS-PAGE. After transfer to PVDF membrane, HRP-conjugated secondary antibody to adiponectin (Table S1) was visualized with Western Lightning Plus (Perkin Elmer, Waltham, Massachusetts) and imaged using a LI-COR Odyssey Imager (LI-COR Biosciences, Lincoln, NE, USA). After immunoblotting, the membrane was stained for 1-minute with Ponceau S as a loading control (0.5% w/v in 1% acetic acid, Fisher, BP103-10). Ponceau stained membranes were rinsed with water prior to drying and imaging.

### Histology and Immunostaining

#### Paraffin immunostaining and imaging

Paraffin embedding, slide preparation, and H&E stains were performed by the WUSM Musculoskeletal Histology and Morphometry core. Bones were fully decalcified in 14% EDTA (Sigma-Aldrich E5134), pH 7.4 prior to embedding. For immunostaining, 10 μm paraffin sections were rehydrated in a series of xylene and ethanols prior to antigen retrieval with 10 mM sodium citrate buffer (pH 6.0, 20-minutes, 90-95°C or overnight at 55°C). Antibodies used for paraffin immunostaining are detailed in Table S1. <u>Paraffin Immunofluorescence:</u> Retrieved sections were permeabilized for 10-minutes in 0.2% Triton-X in PBS, blocked for 1-hour with 10% donkey serum (Sigma-Aldrich D9663) in TNT buffer (0.1 M Tris-HCL pH 7.4, 0.15 M sodium chloride, 0.05% Tween-20), and incubated for 24-hr at 4°C with primary antibodies followed by washing and secondary detection (Table S1). Secondary antibodies in TNT buffer were applied for 1-hour at room temperature. Nuclei were counterstained in 1 μg/mL DAPI (Sigma-Aldrich D9542) for 5-min prior to mounting in Fluoromount-G (ThermoFisher, 00-4958-02). All washes between steps were performed three times each in TNT buffer. <u>Paraffin Immunohistochemistry:</u> Tissue sections were permeabilized for 10-minutes in 0.2% Triton-X in PBS, blocked for 1-hr in kit-specific blocking reagent (ImmPRESS HRP Goat Anti-Rabbit IgG Polymer Detection Kit, Vector Laboratories, MP-7451), and incubated for 24-hr at 4°C with primary antibody (Table S1). Sections were washed in TNT and endogenous peroxidase activity was quenched in 0.3% hydrogen peroxide (Sigma-Aldrich 216763) in PBS for 30-minutes. Sections were then incubated with ImmPRESS polymer reagent for 30-minutes prior to development with peroxidase substrate solution. Slides were counterstained with hematoxylin (Ricca Chemical 3536-16) and dehydrated through a reverse ethanol gradient prior to mounting in Permount. Images were taken using a Nikon Spinning Disk confocal microscope or a Hamamatsu 2.0-HT NanoZoomer System with NDP.scan 2.5 image software.

#### Frozen immunostaining and imaging

Tissues were embedded in OCT mounting media (Fisher HealthCare 23-730-571) and cut at 50 μm on a cryostat (Leica). Bones were fully decalcified in 14% EDTA, pH 7.4 prior to embedding. Sections were blocked in 10% donkey serum in TNT buffer prior to incubation for 48-h with primary antibodies (Table S1). After washing, secondary antibodies in TNT buffer were applied for 24-hours at 4°C (Table S1). The sections were then washed and incubated in DAPI for 5-min prior to mounting with Fluoromount-G. Images were taken at 10x on a Nikon spinning disk confocal microscope.

### Bone marrow stromal cell (BMSC) isolation, cell culture, and immunostaining

Immediately after euthanasia by CO2, long bones were harvested under aseptic conditions. The ends of the long bones were cut to allow flushing of marrow contents, as described previously (37). Cells were suspended in MesenCult Expansion Medium (STEMCELL Technologies 05513) containing MesenCult Basal Medium, MesenCult 1X Supplement, 0.5 mL MesenPure, 1X L-Glutamine, and 1X penicillin/streptomycin. Primary bone marrow cultures were plated at a density of 2.0 x 10^6^ cells/cm^2^ and incubated at 37°C, 5% CO2. After 48-hr, nonadherent cells were removed with subsequent media changes occurring every 2-3 days. After 14 days, colonies were fixed with methanol prior to permeabilization with 1% Triton X-100 (Sigma-Aldrich 9002-93-1) in PBS for 10-min at room temperature. Cells were blocked with a solution containing PBS, 10% donkey serum, and 0.1% Triton X-100 for 30-min prior to incubation for 24-hr at 4°C with primary antibodies (Table S1). Cells were then washed prior to application of secondary antibodies in PBS and 0.1% Triton X-100 for 30-min at room temperature. Nuclei were stained with 1 μg/mL DAPI for 5-min prior to mounting in Fluoromount-G. Images were taken at 4x and 20x using a Nikon spinning disk confocal microscope.

**Table S1.** Antibodies used for western blot and immunostaining.

| Table S1. Antibodies used for western blot and immunostaining. |  |  |  |  |
| --- | --- | --- | --- | --- |
| Western Blot |  |  |  |  |
| Primary Antibody<br>(Vendor, Cat. No) | Dilution | Secondary Antibody<br>(Vendor, Cat. No) | Conjugate | Dilution |
| Rabbit polyclonal Anti-Adiponectin, (Sigma-Aldrich, A6354) | 1:1000 | Anti-Rabbit | HRP | 1:10,000 |
| Paraffin IHC |  |  |  |  |
| Primary Antibody<br>(Vendor, Cat. No) | Dilution | Secondary Antibody<br>(Vendor, Cat. No) | Conjugate | Dilution |
| Anti-CD68<br>(Abcam, UK, ab125212) | 1:1000 | ImmPRESS Reagents<br>(Vector Labs, MP-7401) | HRP | N/A |
| Paraffin Immunofluorescence |  |  |  |  |
| Primary Antibody<br>(Vendor, Cat. No) | Dilution | Secondary Antibody<br>(Vendor, Cat. No) | Fluorophore | Dilution |
| Anti-Perilipin<br>(Progen Biotechnik, Germany, GP29) | 1:400 | Donkey Anti-Guinea Pig<br>(Jackson IR, USA, 706-605-148) | AF647 | 1:200 |
| Frozen Immunofluorescence |  |  |  |  |
| Primary Antibody (Vendor, Cat. No) | Dilution | Secondary Antibody | Fluorophore | Dilution |
| Anti-GFP<br>(Abcam, UK, ab13970) | 1:1000 | Donkey Anti-Chicken<br>(Jackson IR, USA, 703-545-155) | AF488 | 1:500 |
| Anti-RFP<br>(Abcam, UK, ab62341) | 1:500 | Donkey Anti-Rabbit | AF594 | 1:500 |
|  |  | (Jackson IR, USA, 711-585-152) |  |  |
| Anti-Perilipin<br>(Progen Biotechnik, Germany, GP29) | 1:500 | Donkey Anti-Guinea Pig<br>(Jackson IR, USA, 706-605-148) | AF647 | 1:500 |
| **ICC-IF** |  |  |  |  |
| Primary Antibody (Vendor, Cat. No) | Dilution | Secondary Antibody | Fluorophore | Dilution |
| Anti-GFP<br>(Abcam, UK, ab13970) | 1:1000 | Donkey Anti-Chicken<br>(Jackson IR, USA, 703-545-155) | AF488 | 1:500 |
| Anti-RFP<br>(Abcam, UK, ab62341) | 1:500 | Donkey Anti-Rabbit<br>(Jackson IR, USA, 711-585-152) | AF594 | 1:500 |
| Anti-Perilipin<br>(Progen Biotechnik, Germany, GP29) | 1:500 | Donkey Anti-Guinea Pig<br>(Jackson IR, USA, 706-605-148) | AF647 | 1:500 |

### Computed tomography and osmium staining

Bones were embedded in 2% agarose prior to scanning at 20 μm voxel resolution using a Scanco μCT 40 (Scanco Medical AG). Analysis was performed according to reported guidelines (38). For cancellous bone, 100 slices (2 mm) below the growth plate, beginning where the primary spongiosa was no longer visible, were contoured and analyzed at a threshold of 175 (on a 0-1000 scale relative to a pre-calibrated hydroxyapatite phantom). For cortical bone, 20 slices (400 μm) located 2 mm proximal to the tibia-fibula junction were contoured and analyzed at a threshold of 260. To assess bone marrow adiposity, bones were decalcified in 14% EDTA, pH 7.4 and incubated in a solution containing 1% osmium tetroxide (Electron Microscopy Sciences 19170) and 2.5% potassium dichromate (Sigma-Aldrich 24-4520) for 48-hours (39). After washing for 2-hours in running water and storage in PBS at 4°C, osmium-stained bones were embedded in 2% agarose and scanned at 10 μm voxel resolution (Scanco μCT 40; 70 kVp, 114 μA, 300 ms integration time). Regions of interest were contoured for BMAT quantification as detailed in the figure legends. BMAT was segmented with a threshold of 400.

### Bone marrow adipocyte cell size analysis

Tiled 10x images covering the femoral and tibial metaphyses were exported from the Nanozoomer scans of H&E stained slides and processed in Fiji to estimate average adipocyte cell size (40). Based on previous recommendations for adipocyte cell size analyses (41), a minimum of 100 adipocytes were analyzed for each mouse. Briefly, the scale in Fiji was set to be consistent with the original scan. The image was then converted to 8-bit and a threshold of 230 to 255 was applied to create a mask. Then the image was cleaned up using the wand tool and the deletion command to eliminate non-adipocyte structures. The cleaned mask was processed using the Fill Holes and the Watershed tools. The size of adipocytes was determined using the “Analyze Particles” tool by setting the size to 200 to 4000 μm^2^ and circularity to 0.40 – 1.00. Histograms were created in GraphPad Prism and the average adipocyte cell size was calculated using Excel.

### CL316,243 injection

CL316,243 (Sigma-Aldrich, C5976) was reconstituted in saline to a concentration of 0.01 mg/mL and stored at 4°C for up to 2-weeks. Eight daily subcutaneous injections of 0.03 mg/kg CL316,243 were administered over the course of 10-days (weekdays only, M→F, M→W) prior to sacrifice on Day 11.

### Bone marrow adipocyte purification, RNA extraction and qPCR

Bone marrow adipocytes were collected from groups of 8-12 mice at 4- to 6-months of age. Femurs and tibiae (16 to 24 bones/preparation) were rapidly dissected into pre-warmed 37°C HBSS buffer (Gibco 10425-076) containing 2% DNase and protease free bovine serum albumin (Fisher BP9706), 5 mM EDTA, and 1 g/L glucose. After cutting the ends of the bones, whole bone marrow was flushed into a 50 mL conical tube with a 10 mL syringe + 22 gauge needle and resuspended into 20 mL fresh buffer + 1 mg/mL collagenase. Marrow depleted bones were placed into a separate tube in 20 mL buffer + 1 mg/mL collagenase and finely minced to liberate any residual BMAs. Bone and bone marrow preparations were centrifuged at room temperature, 400g x 2 min and BMA-containing supernatant was decanted into a new tube prior to re-centrifugation at 400g x 1 min. Infranatant and any residual pellet was removed using a pulled glass pipet until only 1-2 mL of liquid was remaining. The adipocyte-containing liquid was serially applied to a NucleoSpin^®^ Filter Column (NucleoSpin RNA XS Kit, Takara, 740902) for on-column BMA lysis and RNA extraction. Briefly, the filter column was centrifuged slowly at 50 g x 10 seconds to retain the BMA cells while removing any residual liquid into the collection tube. The bottom of the column was then sealed with parafilm and kit-supplied RNA lysis buffer was added with gentle agitation. BMA-enriched (‘BMAe’) lysates were processed for RNA extraction using the kit-supplied protocol and reagents.

For qPCR, 100 ng of total RNA was reverse transcribed into cDNA using SuperScript IV VILO Master Mix with ezDNase™ Enzyme (Thermo Fisher Scientific 11766050) according to the manufacturer’s instruction. SyGreen 2x Mix Lo-ROX (PCR Biosystems PB20.11-51) was used to perform the qPCR assay on a QuantStudio™ 3 Real-Time PCR System (Thermo Fisher Scientific A28136). Gene expression of individual targets was calculated based on amplification of a standard curve for each primer. Results were normalized to the geometric mean of housekeeping genes *Ppia* and *Tbp*. Primer sequences are listed in Table S2.

**Table S2.**
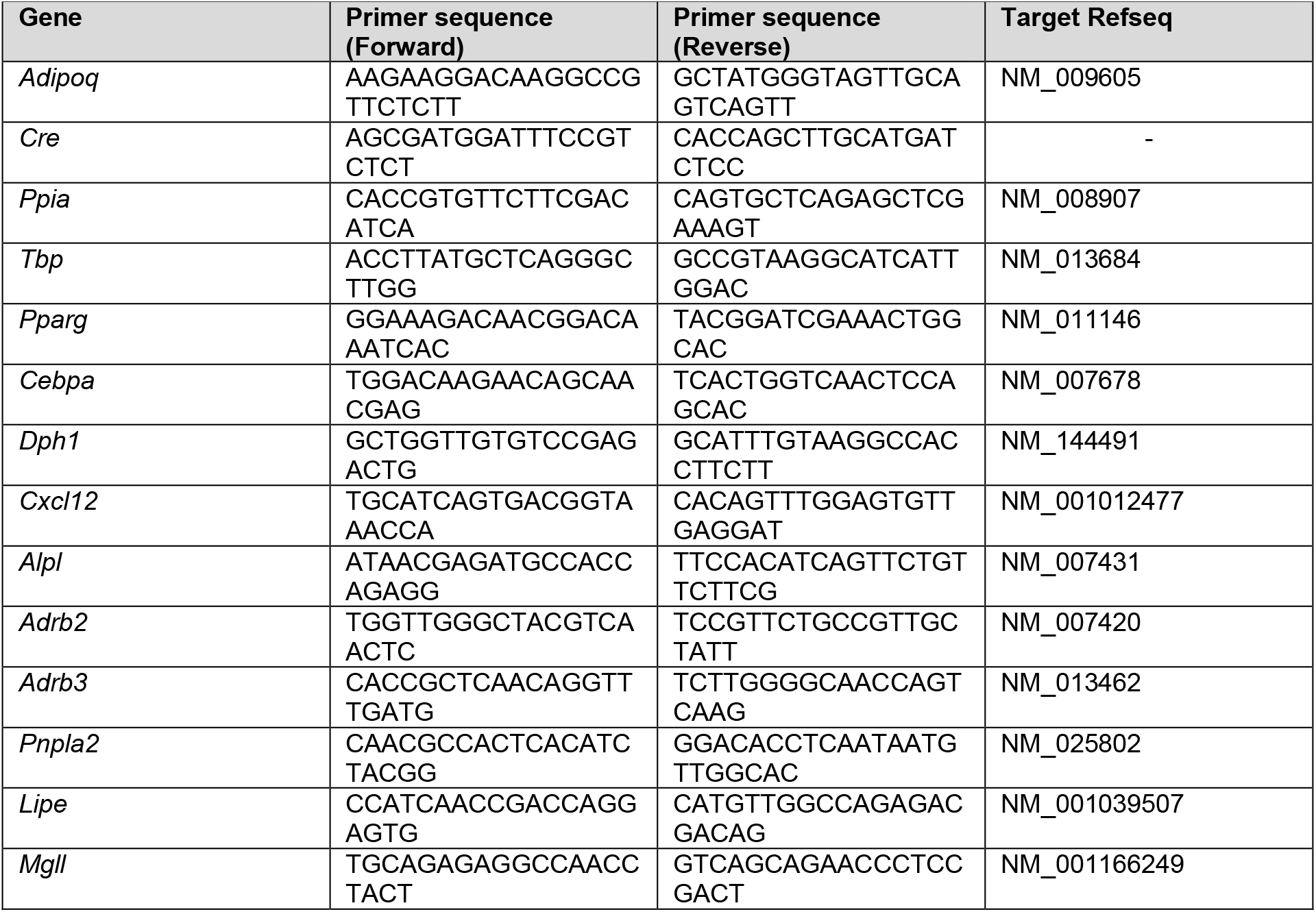
qPCR Primers

### Fat transplantation

*Adipoq*^Cre-/DTA+^ and *Adipoq*^Cre+/DTA+^ mice received subcutaneous fat transplant or sham surgery at 3- to 5-weeks of age. Mice were maintained for an additional 12-weeks prior to sacrifice (end age 15- to 17-weeks). Donor preparation: wild type donor mice on the same background (C57BL/6J) ranged from 22- to 39-days of age. Immediately after decapitation under anesthesia, bilateral inguinal WAT depots were dissected free of surrounding tissues and placed into sterile PBS in a petri dish. The lymph node was removed and a scalpel blade was used to mince the remaining iWAT into small pieces of ~0.5-1.0 mm^3^. The entire minced iWAT from one donor mouse was transplanted to one recipient mouse. Recipient surgery: the recipient mouse was anesthetized with isoflurane and the skin on the back was prepared (shaved and treated 2x each with 70% ethanol and betadine) prior to making two 1 cm incisions along the midline, one over the shoulder blades and one just above the level of the pelvis. Blunt dissection was used to create four pockets just lateral to each incision, one on each side. The minced iWAT from the donor mouse was evenly distributed into the 4 pockets. The incisions were closed and all mice received Buprenex SR at the time of surgery for post-operative analgesia. Post-surgical monitoring and management were performed per DCM guidelines, as approved in our animal protocol.

### Serum glycerol and triglyceride assay

Serum glycerol and true triglyceride (TG) levels were determined using a Serum Triglyceride Determination Kit (Sigma-Aldrich TR0100). In brief, free glycerol reagent and triglyceride reagent were prepared according to the manufacturer’s instruction. To measure serum glycerol, 10 μL serum/well was added to a 96-well microplate on ice prior to addition of 150 μL of free glycerol reagent and incubation at 37°C for 10-minutes. The absorbances of the standards and the samples at 540 nm versus blank (pure Free Glycerol Reagent) were measured using a microplate spectrophotometer (BioTek). To determine serum true TG level, 38 μL of Triglyceride Reagent was added to each well after the initial absorbance measurement for glycerol, followed by an additional 10-minute incubation at 37°C. The absorbances of the standards and the samples at 540 nm versus blank were measured again using the microplate reader. A standard curve was utilized for the calculation of serum-free glycerol and total TG concentrations. The serum true TG level was calculated by subtracting the free glycerol level from the total TG level for each sample, as per manufacturer instructions. All samples were assayed in duplicate.

### Statistics

Statistical analyses were performed in GraphPad Prism including unpaired t-test, one-way, two-way, and three-way ANOVA with multiple comparisons tests, applied as detailed in the figure legends. A p-value of less than 0.05 was considered statistically significant. Quantitative assessments of cell size and μCT-based analyses were performed by individuals that were blinded to the sample identity.

## DATA AVAILABILITY

All relevant data are available from the authors upon reasonable request.

