## Supplemental Figures and Legends for "A novel skeletal-specific adipogenesis pathway defines key origins and adaptations of bone marrow adipocytes with age and disease"

#### SUPPLEMENTAL FIGURE TITLES AND LEGENDS

**Figure S1. Cortical parameters are increased in young *Adipoq*$^{Cre+/DTA+}$ fat free (FF) mice, but tends to decline with age. (A)** Representative *ex vivo* micro-CT cross sections of 4-month old *Adipoq*$^{Cre+/DTA+}$ fat free (FF) and *Adipoq*$^{Cre-/DTA+}$ control (Con) cortical bone at 6 mm, 8 mm, and 12 mm distal to the proximal end of the tibia. **(B-H)** Quantification of cortical bone parameters. **(B)** Cortical thickness. **(C)** Cortical bone volume fraction (Ct. BVF). **(D)** Cortical medullary area. **(E)** Cortical total area (medullary + bone area). **(F)** Cortical bone area. **(G)** Cortical bone mineral content (Ct. BMC). **(H)** Polar moment of inertia (pMOI). Sample size for control and FF mice, respectively: 4-months Male n = 5, 6, Female n = 8, 5; and 8-months Male n = 4, 5; Female n = 3, 3. Statistical significance was assessed by two-way ANOVA with Tukey's multiple comparisons test. ANOVA results as indicated. *p≤0.05. Data presented as mean ± SD. WT and FF mice were housed at 30$^{o}$C on a 12h/12h light/dark cycle.

**Figure S2. Extra-skeletal adipocytes are depleted in the foot and tail of FF mice.** Representative images from *Adipoq*$^{Cre+/DTA+}$ fat free (FF) and *Adipoq*$^{Cre-/DTA+}$ control (Control) mice at 8-months of age. Representative longitudinal hematoxylin and eosin (H&E) stained sections of the **(A)** foot and **(B)** tail vertebrae, including the bone (B) and surrounding soft tissues. Ad = adipocytes; BMA = bone marrow adipocyte. Scale = 0.5 mm. T and FF mice were housed at 30$^{o}$C on a 12h/12h light/dark cycle.

**Figure S3. Engrafted fat transplant fragments resemble subcutaneous white adipose tissue at the time of sacrifice.** Male and female *Adipoq*$^{Cre+/DTA+}$ fat free and *Adipoq*$^{Cre-/DTA+}$ controls underwent sham surgery or were transplanted subcutaneously with control adipose tissue at 3- to 5-weeks of age. After surgery, mice were monitored for 12-weeks prior to sacrifice. At the end point, the transplanted fat engrafted, became vascularized (arrowhead), and grossly resembled subcutaneous white adipose tissue in the FF mice (representative example pictured here).

**Figure S4. Trabecular and cortical bone phenotypes remain unchanged after subcutaneous fat transplant.** Male and female *Adipoq*$^{Cre+/DTA+}$ fat free (FF) and *Adipoq*$^{Cre-/DTA+}$ control (Con) mice underwent sham surgery or were transplanted subcutaneously with WT adipose tissue at 3- to 5-weeks of age. After surgery, mice were monitored for 12-weeks prior to sacrifice. **(A)** Representative µCT images of tibiae from male and female mice, as indicated. Scale = 1 mm. **(B)** Cortical thickness. **(C)** Cortical medullary area. **(D)** Trabecular bone volume fraction (Tb. BVF). **(E)** Trabecular bone number (Tb. Number). Sample size for control and FF mice, respectively: sham Male n = 4, 6; transplant Male n = 4, 5; sham Female n = 5, 7; transplant Female n = 5, 4. Statistical significance was assessed by two-way ANOVA with Tukey's multiple comparisons test. ANOVA results as indicated. *p≤0.05. Data presented as mean ± SD. All mice were housed at 30$^{o}$C on a 12h/12h light/dark cycle.

**Figure S5. Adipose tissue masses remain unchanged after subcutaneous fat transplant.** Male and female *Adipoq*$^{Cre+/DTA+}$ fat free (FF) and *Adipoq*$^{Cre-/DTA+}$ control (Con) mice underwent sham surgery or were transplanted subcutaneously with WT adipose tissue at 3- to 5-weeks of age. After surgery, mice were monitored for 12-weeks prior to sacrifice. Endpoint tissue weights for *Adipoq*$^{Cre+/DTA+}$ mice and *Adipoq*$^{Cre-/DTA+}$ **(A)** brown adipose tissue (BAT), **(B)** inguinal white adipose tissue (iWAT), and **(C)** gonadal adipose tissue (gWAT). Sample size for control and FF mice, respectively: sham Male n = 4, 6; transplant Male n = 4, 5; sham Female n = 5, 7; transplant Female n = 5, 4. Statistical significance was assessed by two-way ANOVA with Tukey's multiple comparisons test. ANOVA results as indicated. *p≤0.05. Data presented as mean ± SD. All mice were housed at 30$^{o}$C on a 12h/12h light/dark cycle.

### Figure S1

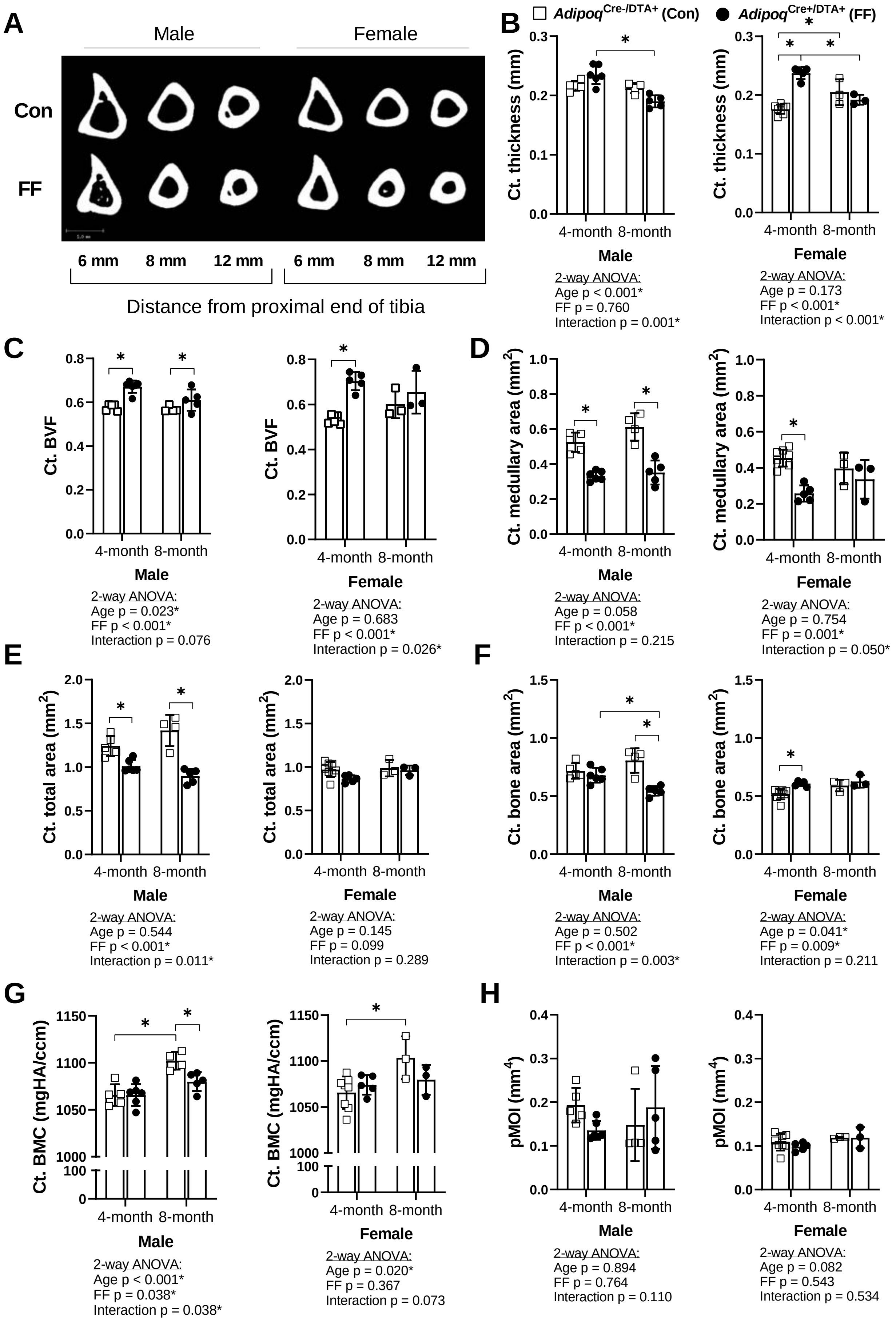
A
Male
Female
Con
FF
6 mm
8 mm
12 mm
6 mm
8 mm
12 mm
Distance from proximal end of tibia
B
Adipoq Cre-/DTA+ (Con)
Adipoq Cre+/DTA+ (FF)
Ct. thickness (mm)
4-month 8-month
Male
2-way ANOVA:
Age p < 0.001*
FF p = 0.760
Interaction p = 0.001*
Female
2-way ANOVA:
Age p = 0.173
FF p < 0.001*
Interaction p < 0.001*
C
Ct. BVF
Male
2-way ANOVA:
Age p = 0.023*
FF p < 0.001*
Interaction p = 0.076
Female
2-way ANOVA:
Age p = 0.683
FF p < 0.001*
Interaction p = 0.026*
D
Ct. medullary area (mm2)
Male
2-way ANOVA:
Age p = 0.058
FF p < 0.001*
Interaction p = 0.215
Female
2-way ANOVA:
Age p = 0.754
FF p = 0.001*
Interaction p = 0.050*
E
Ct. total area (mm2)
Male
2-way ANOVA:
Age p = 0.544
FF p < 0.001*
Interaction p = 0.011*
Female
2-way ANOVA:
Age p = 0.145
FF p = 0.099
Interaction p = 0.289
F
Ct. bone area (mm2)
Male
2-way ANOVA:
Age p = 0.502
FF p < 0.001*
Interaction p = 0.003*
Female
2-way ANOVA:
Age p = 0.041*
FF p = 0.009*
Interaction p = 0.211
G
Ct. BMC (mgHA/ccm)
Male
2-way ANOVA:
Age p < 0.001*
FF p = 0.038*
Interaction p = 0.038*
Female
2-way ANOVA:
Age p = 0.020*
FF p = 0.367
Interaction p = 0.073
H
pMOI (mm4)
Male
2-way ANOVA:
Age p = 0.894
FF p = 0.764
Interaction p = 0.110
Female
2-way ANOVA:
Age p = 0.082
FF p = 0.543
Interaction p = 0.534

**Figure S2**

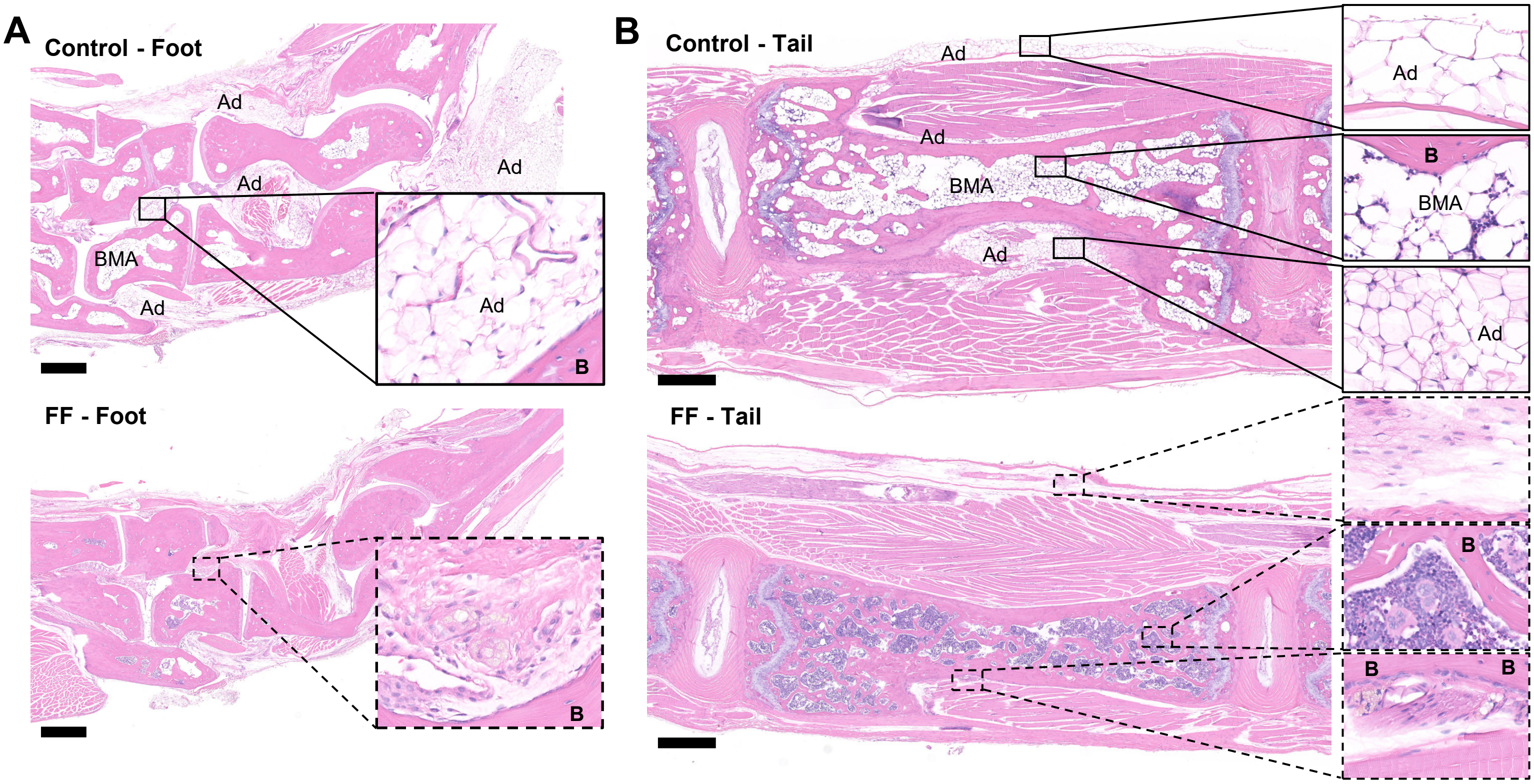

A
Control - Foot
Ad
Ad
Ad
BMA
Ad
Ad
B
FF - Foot
B
B
Control - Tail
Ad
Ad
BMA
Ad
Ad
B
BMA
Ad
FF - Tail
B
B
B

**Figure S3**

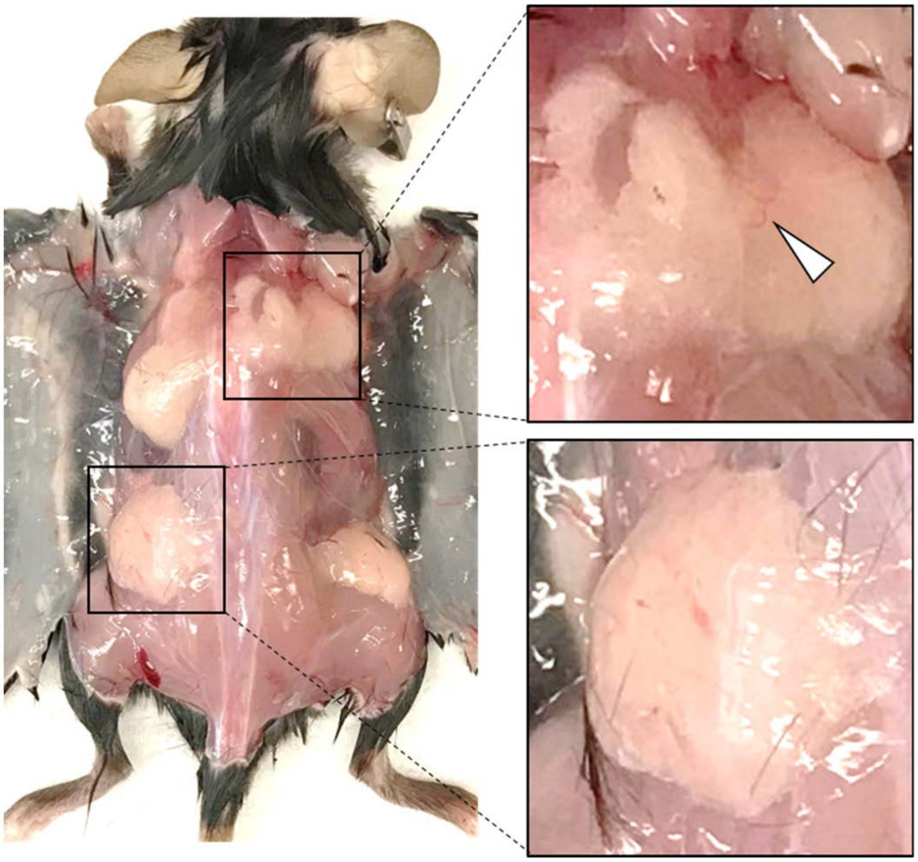

### Figure S4

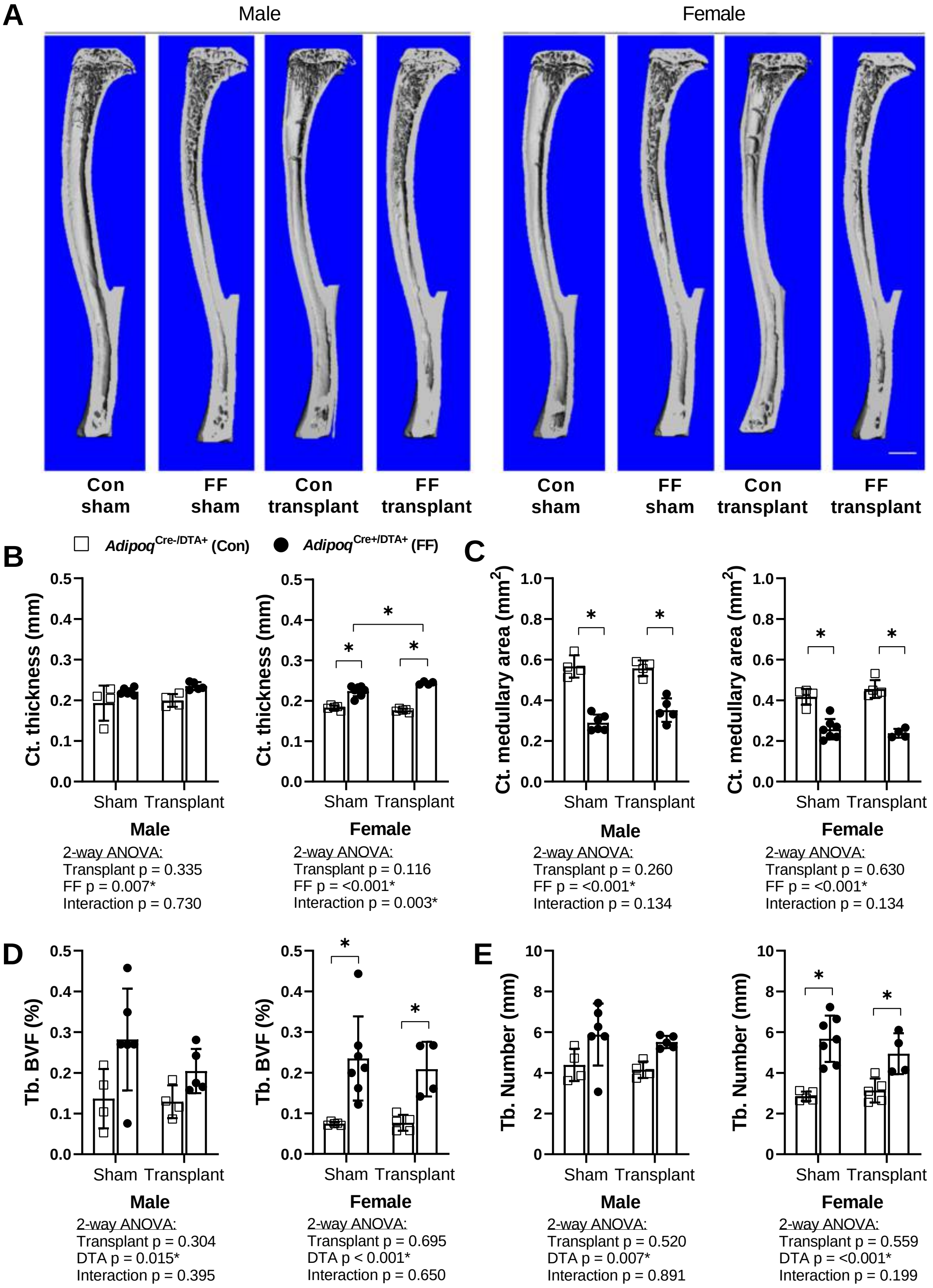
A
Male
Female
Con sham
FF sham
Con transplant
FF transplant
Con sham
FF sham
Con transplant
FF transplant
B
C
Adipoq Cre-/DTA+ (Con)
Adipoq Cre+/DTA+ (FF)
Ct. thickness (mm)
Ct. medullary area (mm2)
Sham Transplant
Male
Female
2-way ANOVA:
Transplant p = 0.335
FF p = 0.007*
Interaction p = 0.730
2-way ANOVA:
Transplant p = 0.116
FF p = <0.001*
Interaction p = 0.003*
2-way ANOVA:
Transplant p = 0.260
FF p = <0.001*
Interaction p = 0.134
2-way ANOVA:
Transplant p = 0.630
FF p = <0.001*
Interaction p = 0.134
D
E
Tb. BVF (%)
Tb. Number (mm)
2-way ANOVA:
Transplant p = 0.304
DTA p = 0.015*
Interaction p = 0.395
2-way ANOVA:
Transplant p = 0.695
DTA p < 0.001*
Interaction p = 0.650
2-way ANOVA:
Transplant p = 0.520
DTA p = 0.007*
Interaction p = 0.891
2-way ANOVA:
Transplant p = 0.559
DTA p = <0.001*
Interaction p = 0.199

### Figure S5

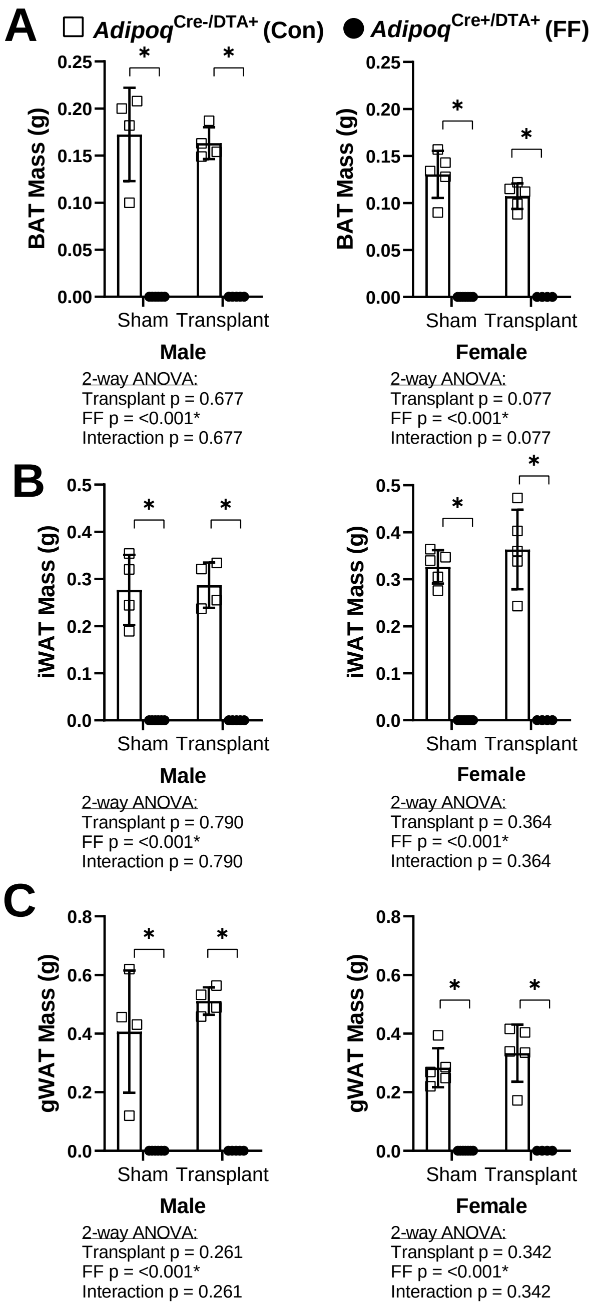
A
Adipoq Cre-/DTA+ (Con)
Adipoq Cre+/DTA+ (FF)
BAT Mass (g)
0.25
0.20
0.15
0.10
0.05
0.00
*
Sham
Transplant
Male
2-way ANOVA:
Transplant p = 0.677
FF p = <0.001*
Interaction p = 0.677
Female
2-way ANOVA:
Transplant p = 0.077
FF p = <0.001*
Interaction p = 0.077
B
iWAT Mass (g)
0.5
0.4
0.3
0.2
0.1
0.0
Male
2-way ANOVA:
Transplant p = 0.790
FF p = <0.001*
Interaction p = 0.790
Female
2-way ANOVA:
Transplant p = 0.364
FF p = <0.001*
Interaction p = 0.364
C
gWAT Mass (g)
0.8
0.6
0.4
0.2
0.0
Male
2-way ANOVA:
Transplant p = 0.261
FF p = <0.001*
Interaction p = 0.261
Female
2-way ANOVA:
Transplant p = 0.342
FF p = <0.001*
Interaction p = 0.342
